# Yolk granule fusion and microtubule aster formation regulate cortical granule translocation and exocytosis in zebrafish oocytes

**DOI:** 10.1101/2022.08.10.503442

**Authors:** Shayan Shamipour, Laura Hofmann, Irene Steccari, Roland Kardos, Carl-Philipp Heisenberg

## Abstract

Dynamic reorganization of the cytoplasm is key to many core cellular processes, such as cell division, cell migration and cell polarization. Cytoskeletal rearrangements are thought to constitute the main drivers of cytoplasmic flows and reorganization. In contrast, remarkably little is known about how dynamic changes in size and shape of cell organelles affect large-scale cytoplasmic organization. Here, we show that within the maturing zebrafish oocyte, the surface localization of exocytosis-competent cortical granules upon germinal vesicle breakdown is achieved by the combined activities of yolk granule fusion and microtubule aster formation and translocation. We find that cortical granules are moved towards the oocyte surface through radially-outward cytoplasmic flows induced by yolk granules fusing within the oocyte center in response to GV breakdown. We further show that vesicles decorated with the small Rab GTPase Rab11, a master regulator of vesicular trafficking and exocytosis, accumulate together with cortical granules at the oocyte surface. This accumulation is achieved by Rab11-positive vesicles being transported by acentrosomal microtubule asters, the formation of which is induced by the release of CyclinB/Cdk1 upon GV breakdown, and which display a net movement towards the oocyte surface by preferentially binding to the oocyte actin cortex. We finally demonstrate that the decoration of cortical granules by Rab11 at the oocyte surface is needed for cortical granule release and subsequent chorion elevation, a process central in oocyte activation. Collectively, these findings unravel a yet unrecognized role of organelle fusion, functioning together with cytoskeletal rearrangements, in determining cytoplasmic organization during oocyte maturation.

## Introduction

Oogenesis marks the very first step in development, establishing the maternal blueprint for embryonic patterning. During this process, the oocyte grows in size by acquiring maternally-provided material and completes its first meiosis to eventually become arrested in the metaphase of meiosis-II until fertilization occurs (Wallace and Selman, 1981). Central to oogenesis is the accurate positioning of large organelles, such as the oocyte nucleus (Germinal vesicle, GV) and meiotic spindle, but also small fate-determining molecules, such as mRNAs and proteins, within the oocyte, a process fundamental for embryonic axis formation and cell fate specification (Almonacid et al., 2015; Azoury et al., 2008; Lénárt et al., 2005; Prodon et al., 2008, 2006; Quinlan, 2016; Schuh and Ellenberg, 2008; Yi et al., 2013). Yet, how such positioning of ooplasmic components is orchestrated in space and time is still only poorly understood.

Cytoplasmic organization can occur in the absence of external cues, suggesting that the cytoplasm is capable of self-organization (Mitchison and Field, 2021; Shamipour et al., 2021). Previous research has highlighted an important role for the cell cytoskeleton, and especially the microtubule and actin networks, in driving such cytoplasmic self-organization (Cheng and Ferrell, 2019; Ierushalmi et al., 2020; Sakamoto et al., 2020). For instance, microtubules and the movement of motors along microtubule tracks can power cytoplasmic flows and the repositioning of microtubule asters by generating viscous drag forces to the surrounding cytoplasm (Cheng and Ferrell, 2019; Kimura et al., 2017; Meaders et al., 2020; Monteith et al., 2016; Xie and Minc, 2020). Likewise, myosin II-dependent contractions of both cortical and bulk actin networks can result in large-scale actomyosin network flows, which in turn drag the adjacent cytoplasm via friction forces acting at their interface (Deneke et al., 2019; Ierushalmi et al., 2020; Shamipour et al., 2019). In addition to these motor-dependent processes, actin polymerization on the surface of organelles can drive organelle motility, thereby generating active diffusion within the bulk of the cytoplasm (Li et al., 2008; Shamipour et al., 2019; Theriot et al., 1992). However, to what extent cellular processes other than cytoskeletal rearrangements, such as dynamic fusion and splitting of organelles, also function in large-scale cytoplasmic reorganization, remains unclear.

To tackle this question, we turned to the last stage of zebrafish oogenesis (oocyte maturation), during which large-scale ooplasmic reorganizations are accompanied by changes in organelle shape, size and position, preparing the oocyte for fertilization and embryonic development (Selman et al., 1993). Zebrafish oogenesis constitutes a five-stage process: during stages I and II, the oocyte animal-vegetal axis becomes determined through the vegetal pole localization of the Balbiani body, a membrane-less structure rich in mitochondria, proteins and mRNAs required for the vegetal pole establishment (Elkouby et al., 2016). Stages II and III mark the formation of cortical granules (Cgs), which will be exocytosed upon fertilization to prevent polyspermy, and yolk granules (Ygs), which function as energy reservoirs for subsequent embryonic development (Selman et al., 1993). Stage-III oocytes remain arrested in prophase I until oocyte maturation begins. During oocyte maturation (stage-IV), a multitude of transitions take place within the oocyte. For instance, the GV migrates to and breaks down at the animal pole of the oocyte, which is followed by ooplasm accumulating at the animal pole, forming a yolk-free blastodisc, and Cgs obtaining exocytosis-competency, which together give rise to a fertilizable egg (Fernández et al., 2006; Fuentes et al., 2018; Lessman, 2009; Selman et al., 1993). How these different aspects of cytoplasmic reorganization are spatiotemporally orchestrated is still largely unknown.

Here, we show that Cg accumulation at the oocyte surface and acquisition of exocytosis competency upon GV breakdown are driven by the concerted activities of microtubule network rearrangements and Yg fusion. The microtubule network functions in this process by reorganizing into acentrosomal aster-like structures that collectively translocate towards the oocyte surface, taking along Rab11-positive vesicles, while Yg fusion towards the oocyte center induces radially outward cytoplasmic flows that lead to the translocation and accumulation of Cgs at the oocyte surface. Finally, the decoration of Cgs by Rab11 at the oocyte surface confers competency to Cgs to be exocytosed during oocyte activation.

## Results

### GV breakdown triggers blastodisc formation, Yg fusion and Cg outward flow

To unravel the molecular, cellular and biophysical mechanisms underlying ooplasmic reorganization during zebrafish oocyte maturation, we first analyzed how those processes occur in space and time. To this end, ovaries of female zebrafish were harvested, and stage-III oocytes were isolated according to their size and ooplasmic opacity and exposed to the steroid hormone DHP triggering their maturation (Lessman, 2009; Selman et al., 1993). To determine which processes take place during oocyte maturation, we monitored ooplasmic reorganization in oocytes from Tg(*hsp:Clip170-eGFP*) females, in which the ooplasm is labeled by GFP (Fig. 1A and video S1). We found that following GV translocation to and breakdown at the animal pole of the oocyte, ooplasm accumulated at the animal pole, indicative of blastodisc formation (Fig. 1A, S1A and S1A’). In order to visualize which other ooplasmic rearrangements occur upon GV breakdown, we marked both Ygs and Cgs within the ooplasm by exposing Tg(*hsp:Clip170-eGFP*) oocytes to Lysotracker dye, which exclusively labels Ygs but not Cgs, allowing us to distinguish between these different granule types (Fig. 1B and video S1). We further confirmed that granules not labeled by Lysotracker were indeed Cgs by showing that within the mature oocyte/egg they colocalized with Rab11 and underwent exocytosis upon egg activation (Fig. 5C), features typically associated with Cgs (Kanagaraj et al., 2014). Analyzing changes in the subcellular distribution of Ygs and Cgs within the oocyte upon GV breakdown revealed that Ygs were fusing, resulting in a two-fold increase in their average crosssectional area (Fig. 1C–1D and video S1). At the same time, the density of Cgs increased in cortical regions of the oocyte (Fig. 1D and S1B), due to their translocation from central to cortical regions (Fig. 1B–1B’ and video S1). This notion was further supported by tracking individual Cgs, revealing a net movement of Cgs from the center towards the oocyte cortex concomitant with the fusion and compaction of Ygs at the oocyte center (Fig. 1E and video S1). Collectively, these findings suggest that GV breakdown is temporally correlated with blastodisc formation at the AP, Yg fusion and Cg translocation to the oocyte cortex (Fig. S1D).

**Figure 1:**
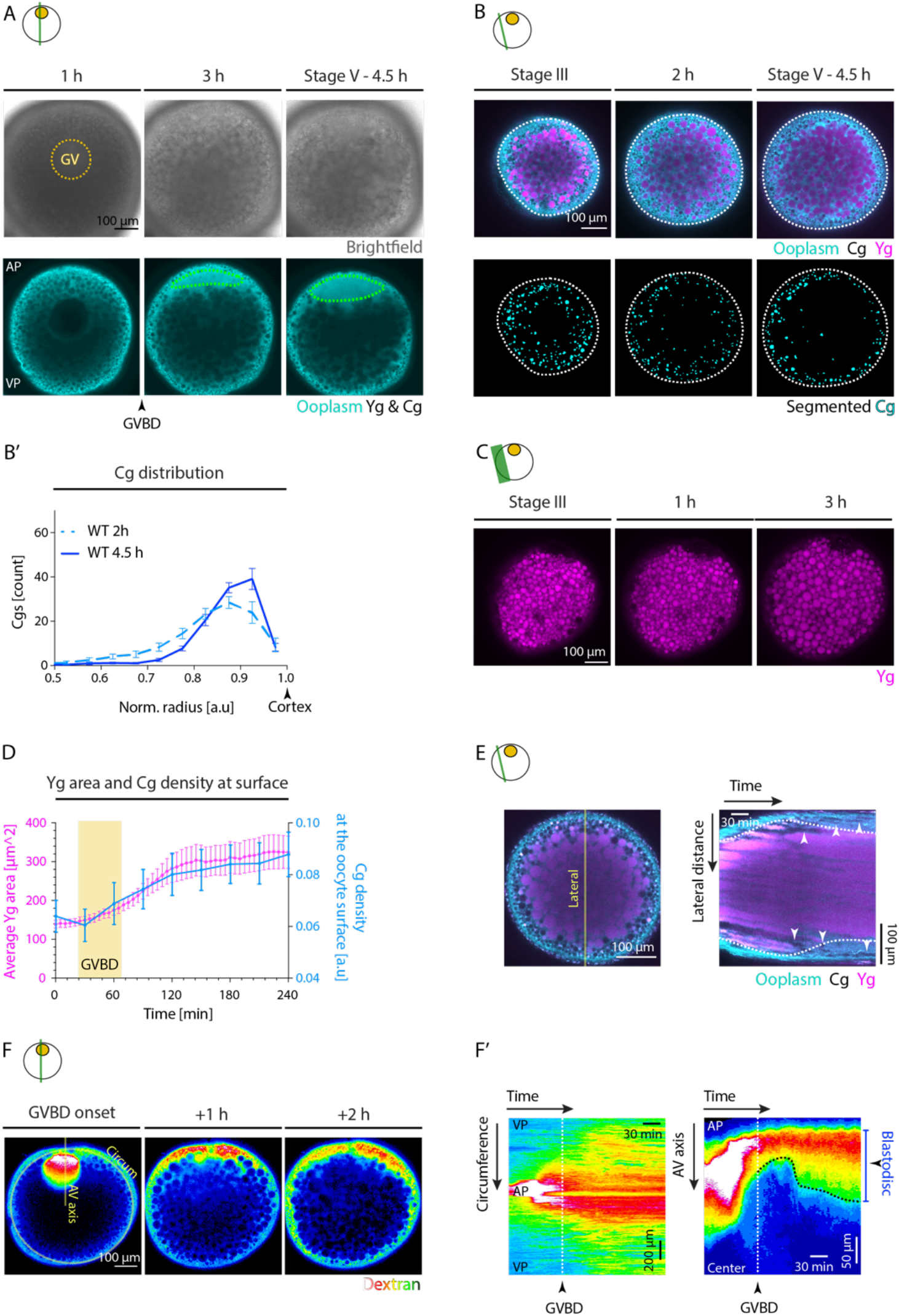
Ooplasmic reorganizations during zebrafish oocyte maturation. (**A**) Bright-field (top row) and fluorescence (bottom row) images of Tg(*hsp:clip170-GFP*) oocytes labeling the ooplasm during oocyte maturation at 1, 3 and 4.5 hours (h) after maturation induction with the DHP hormone. Dashed yellow circle highlights the germinal vesicle (GV) contour and cyan lines indicate the blastodisc region. Arrowhead denotes the germinal vesicle breakdown (GVBD) onset. AP, animal pole; VP, vegetal pole. (**B**) Fluorescence images of stage-III Tg(*hsp:clip170-GFP*) oocytes labeling ooplasm (cyan) and exposed to Lysotracker to mark yolk granules (Yg, magenta) and cortical granules (Cg, black, identified by their exclusion of both Clip-170-GFP and Lysotracker) before maturation onset (stage-III), and 2 and 4.5 h/stage-V after maturation onset (top row). Images in the bottom row show segmented Cg obtained from the images in the top row. Dashed lines mark the oocyte outline. (**B’**) Cg density profile along the oocyte radius at 2 and 4.5 h after maturation onset. Normalized (norm) radius of 0 and 1 correspond to the oocyte center and surface, respectively (N=3 experiments, n=12 oocytes). WT, *wild type* oocytes. (**C**) Maximum fluorescence intensity projection of stage-III oocytes exposed to Lysotracker to label Yg (magenta) before maturation (stage-III) and 1 and 3 h after maturation onset. (**D**) Average Yg area (magenta line, N=3, n=20) and Cg density at the oocyte surface (cyan line, N=3, n=13) over time. Yellow box indicates the period during which GVBD takes place. (**E**) Left: Fluorescence image of Tg(*hsp:clip170-GFP*) oocytes to mark ooplasm (cyan) and exposed to Lysotracker to mark Yg (magenta) and Cg (black, identified by their exclusion of both Clip-170-GFP and Lysotracker). The yellow line along the lateral axis was used to obtain the kymograph shown on the right. Right: Kymograph acquired along the lateral axis of the oocyte shown on the left as a function of time. Arrowheads trace individual Cg moving towards the oocyte surface. (**F**) Fluorescence images of oocytes injected with Dextran Alexa 647 to mark GV nucleoplasm at the onset, and 1 and 2 h post GVBD. Yellow lines along the oocyte circumference (Circum) and animal-vegetal (AV) axis were used to obtain the kymographs in F’. (**F’**) Kymographs acquired along the circumference (left) and AV axis (right) of the oocyte shown on the left as a function of time. The white dashed lines mark the time point of GVBD. The black dashed line tracks the blastodisc interface. Schematics demarcate the imaging plane used for obtaining the images in each panel. Note that in panels (A and F) processes deeper within the oocyte are captured, while in panels (B, C and E) more superficial parts of the oocyte are captured. Error bars, SEM.

### Bulk actomyosin drives blastodisc expansion

Our finding that blastodisc formation is spatiotemporally linked to GV breakdown at the animal pole of the oocyte (Fig. 1A) suggests that these processes might be functionally linked. Given the large size of the GV, accounting for nearly ~ 1.5 % of the total stage-III oocyte volume, we hypothesized that the nucleoplasm released after GV breakdown might directly result in blastodisc formation. To test this possibility, we labeled the nucleoplasm stored within the GV with fluorescently-labeled Dextran and followed its subcellular distribution after GV breakdown (Fig. 1F and video S1). This showed that despite the apparent rapid diffusion of the nucleoplasm from the point of GV breakdown at the animal pole towards the vegetal pole of the oocyte, the majority of the nucleoplasm still remained at the animal pole, thereby initiating blastodisc formation (Fig. 1F, 1F’ and S1C). Moreover, the size of the blastodisc continued to grow after the completion of GV breakdown, rather than shrink as expected for continuous diffusion of the nucleoplasm away from the animal pole (Fig. 1F’), suggesting that mechanisms other than GV nucleoplasm release must be involved in blastodisc formation.

Bulk actomyosin network contraction and flows have previously been implicated in blastodisc expansion within the mature/fertilized oocyte (Shamipour et al., 2019). To determine whether actomyosin network contraction is also involved in the initial blastodisc formation during oocyte maturation, we exposed immature stage-III oocytes to inhibitors specifically interfering with actin polymerization (Cytochalasin B, Cyto B) and myosin-II activity (para-Nitroblebbistatin, PBb), and monitored how such treatment affects blastodisc formation. We found that in oocytes treated with 30 μg/ml Cyto B or 100 μM PBb, blastodisc expansion was strongly diminished (Fig. S2A-A” and video S2), indicating that actomyosin network contraction is required for this process.

To determine how actomyosin network contraction functions in blastodisc formation, we analyzed dynamic changes in the intensity of F-actin during oocyte maturation using Tg(*actb1:Utr-GFP*) oocytes labeling F-actin. This analysis revealed that actin became enriched within the GV and was released to the surrounding ooplasm upon GV breakdown (Fig. 2A-A’ and S3B and video S3), where it diffused away from the animal pole towards the vegetal pole (Fig. S3A-A’). To examine whether these processes establish an actin gradient along the animal-vegetal axis of the oocyte, we visualized F-actin distribution in sections of stage-IV zebrafish oocytes that had just undergone GV breakdown using Phalloidin staining. This revealed an animal-to-vegetal bulk actin gradient with peak levels close to the animal pole of the oocyte (Fig. 2B-B’). Such a bulk actin gradient, analogous to the situation in mature oocytes after fertilization, might lead to bulk actomyosin flows directed towards the animal pole of the oocyte, which, by dragging along the ooplasm, then triggers the accumulation of ooplasm at the animal pole leading to blastodisc formation. Importantly, since the bulk actin gradient was largely limited to the vicinity of the animal pole (Fig. 2B’), we assumed that the resultant flows will also be locally restricted to the animal pole. Consistent with this assumption, measuring ooplasmic flows along the animal-vegetal oocyte axis revealed that they occur predominantly close to the animal pole of the oocyte (Fig. 2C-C’ and video S3), further supporting the notion that bulk actomyosin flows drive ooplasmic flows towards the animal pole, leading to blastodisc formation. Notably, bulk actin levels ceased once the first meiosis was completed (Fig. S3 and video S4), raising questions as to the mechanisms by which the size of blastodisc is maintained during second meiosis where no clear bulk actin was detectable anymore within the oocyte ooplasm (Fig. S3B). Strikingly, we observed actin comet-like structures forming on the surface of Ygs which were located close to the blastodisc interface thereby preventing their return into the blastodisc region (Fig. 2D). This mechanism is highly reminiscent of the function of actin comet-like structures on Ygs in mature oocytes promoting ooplasm-Yg segregation (Shamipour et al., 2019). Together, our results suggest that - similar to the situation in mature oocytes - the combined function of actin flows towards the animal pole and actin comets on the surface of Ygs are responsible for blastodisc formation and maintenance during oocyte maturation (Fig. S3D).

**Figure 2:**
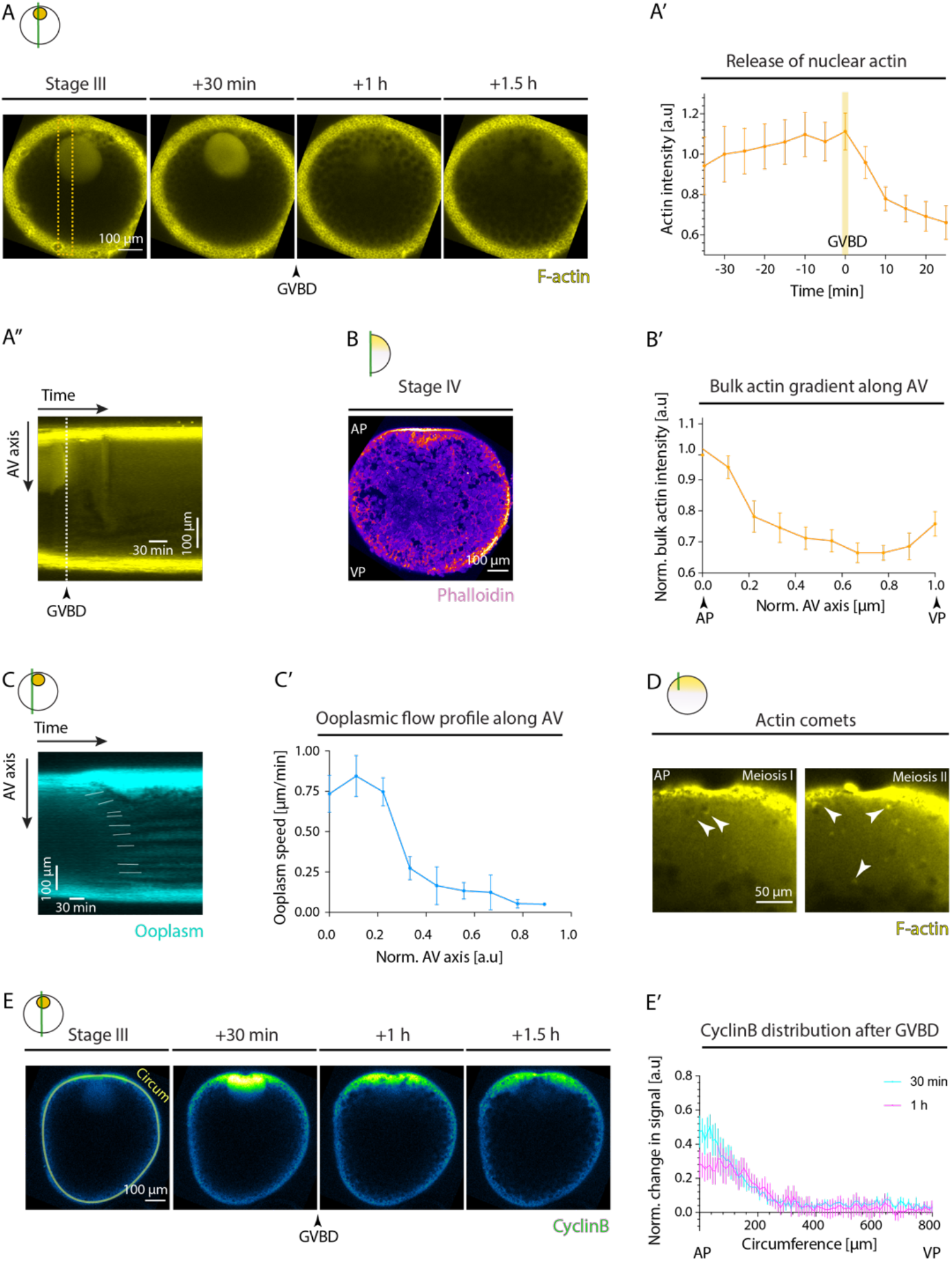
Rearrangement of the actin cytoskeleton during oocyte maturation. (**A**) Fluorescence images of stage-III Tg(*actb1:Utr-GFP*) oocytes labeling F-actin during consecutive stages before maturation (stage-III) and 30 min, 1 h and 1.5 h after maturation induction with the DHP hormone. Arrowhead denotes the germinal vesicle breakdown (GVBD) onset. The dashed box indicates the animal-vegetal (AV) axis used for acquiring the kymograph in (A’’). (**A’**) Normalized bulk actin intensity in the blastodisc region as a function of time (N=2 experiments, n=23 oocytes). Yellow box indicates the GVBD onset. (**A’’**) Kymograph acquired along the animal-vegetal (AV) axis of the oocyte in (A) as a function of time. The white dashed line marks GVBD onset. (**B**) Fluorescence image of a stage-IV oocyte fixed, sectioned and stained with Phalloidin to visualize and measure bulk F-actin gradient along the AV oocyte axis shown in (B’). AP, animal pole; VP, vegetal pole (**B’**) Normalized bulk actin profile along the AV axis, measured from fixed stage-IV oocytes stained with phalloidin as in (B), (N=1, n=10). (**C**) Kymograph acquired along the AV axis of Tg(*hsp:clip170-GFP*) oocytes marking ooplasm as a function of time. White lines trace the flow of ooplasmic pockets over time. (**C’**) Ooplasmic flow profile along the AV axis, measured from the slopes of ooplasmic flow trajectories as shown in (C). Normalized AV of 0 and 1 correspond to the animal and vegetal poles, respectively (N=3, n=17). (**D**) Fluorescence images of oocytes injected with 200 pg *Utrophin-GFP* mRNA to label F-actin during first and second meiosis corresponding to 135 and 170 min after maturation induction with the DHP hormone, respectively. White arrowheads mark actin comets forming within the blastodisc region on the surface of yolk granules. (**E**) Fluorescence images of stage-III oocytes injected with *CyclinB-GFP* mRNA before (stage-III) and 30 min, 1 h and 1.5 h after maturation onset. Arrowhead denotes the GVBD onset. The yellow line along the oocyte circumference (Circum) was used to acquire the intensity profiles plotted in (E’). (**E’**) Normalized change in CyclinB signal relative to its distribution at the time prior to GVBD, measured at 30 min (cyan) and 1 h (magenta) after GVBD along the oocyte circumference (the yellow line in E). Circumference of 0 and 800 μm correspond to the AP and VP, respectively (N=2, n=5). Schematics in each panel demarcate the imaging plane used for obtaining the images in that panel. Error bars, SEM.

Finally, given that the changes in bulk actin dynamics were concomitant with cell cycle progression (Fig. S3), we asked whether CyclinB/Cdk1 complex, the key cell cycle regulator previously suggested to induce bulk actin polymerization and flows within the mature oocyte (Shamipour et al., 2019), might also function as effector by which GV breakdown triggers bulk actin polymerization and flows and consequently blastodisc formation during oocyte maturation. To this end, we injected *CyclinB-GFP* mRNA into stage-III oocytes and visualized its dynamics as a proxy for CyclinB/Cdk1 activity within the maturing oocyte (Bischof et al., 2017). We found CyclinB to be highly enriched within the GV of stage-III oocytes and then released to the surrounding ooplasm upon GV breakdown (Fig. 2E and video S3). Importantly, the release of CyclinB led to a spatially restricted gradient of CyclinB at the oocyte animal pole, which persisted for more than 1 hour after GV breakdown (Fig. 2E’), highly reminiscent of the gradients observed for the nucleoplasm (Fig. 1F-F’ and S1C) and bulk actin (Fig. 2B-B’) following GV breakdown. Moreover, inhibiting CyclinB synthesis by exposing stage-IV oocytes undergoing maturation to 700 μM Cycloheximide resulted in decreased ooplasmic flows and blastodisc formation (Fig. S2A-A” and video S2). This suggests that the activation and release of CyclinB/Cdk1 at the animal pole of the oocyte drives bulk actin polymerization, which in turn triggers bulk actin and ooplasmic flows leading to blastodisc formation.

### Microtubule network forms asters upon GV breakdown

To determine whether other cytoskeletal elements, and in particular microtubules, might also have a function in ooplasmic reorganization during oocyte maturation, we exposed immature stage-III oocytes to inhibitors specifically interfering with microtubule assembly/disassembly (Colchicine and Taxol), and monitored how such treatment affects ooplasmic reorganization and oocyte maturation. We found that in oocytes treated with 200 μM of Colchicine blastodisc formation was largely unaffected (Fig. S2A-A” and video S2), suggesting that microtubules might be dispensable for this process. Interestingly, however, both Colchicine and Taxol-treated oocytes displayed strongly reduced chorion elevation, a process previously shown to be mediated by the release of Cgs at the oocyte surface (Kanagaraj et al., 2014; Fig. S2B-B’). To understand whether and how microtubules might be involved in Cg relocalization and/or exocytosis during oocyte maturation, we first monitored how the microtubule cytoskeleton changes during oocyte maturation in oocytes from Tg(*XlEef1a1:dclk2-GFP*) fish labeling microtubules. We found microtubules to be uniformly distributed throughout the ooplasm of stage-III oocytes (Fig. 3A). This microtubule network, however, transformed drastically upon GV breakdown with numerous bright foci of microtubules appearing in the bulk of the ooplasm (Fig. 3A and video S5). Closer examination of those foci revealed that they resembled microtubule aster-like structures undergoing dynamic fusion and splitting over time (Fig. 3B and video S5), the number of which increased shortly after GV breakdown and then dropped again (Fig. 3C). These consecutive phases of aster formation and disappearance gave rise to a microtubule transformation wave propagating from the animal to the vegetal pole of the oocyte, where asters were forming at the leading edge and dissolving again at the trailing edge (Fig. 3A’ and S4A).

**Figure 3.**
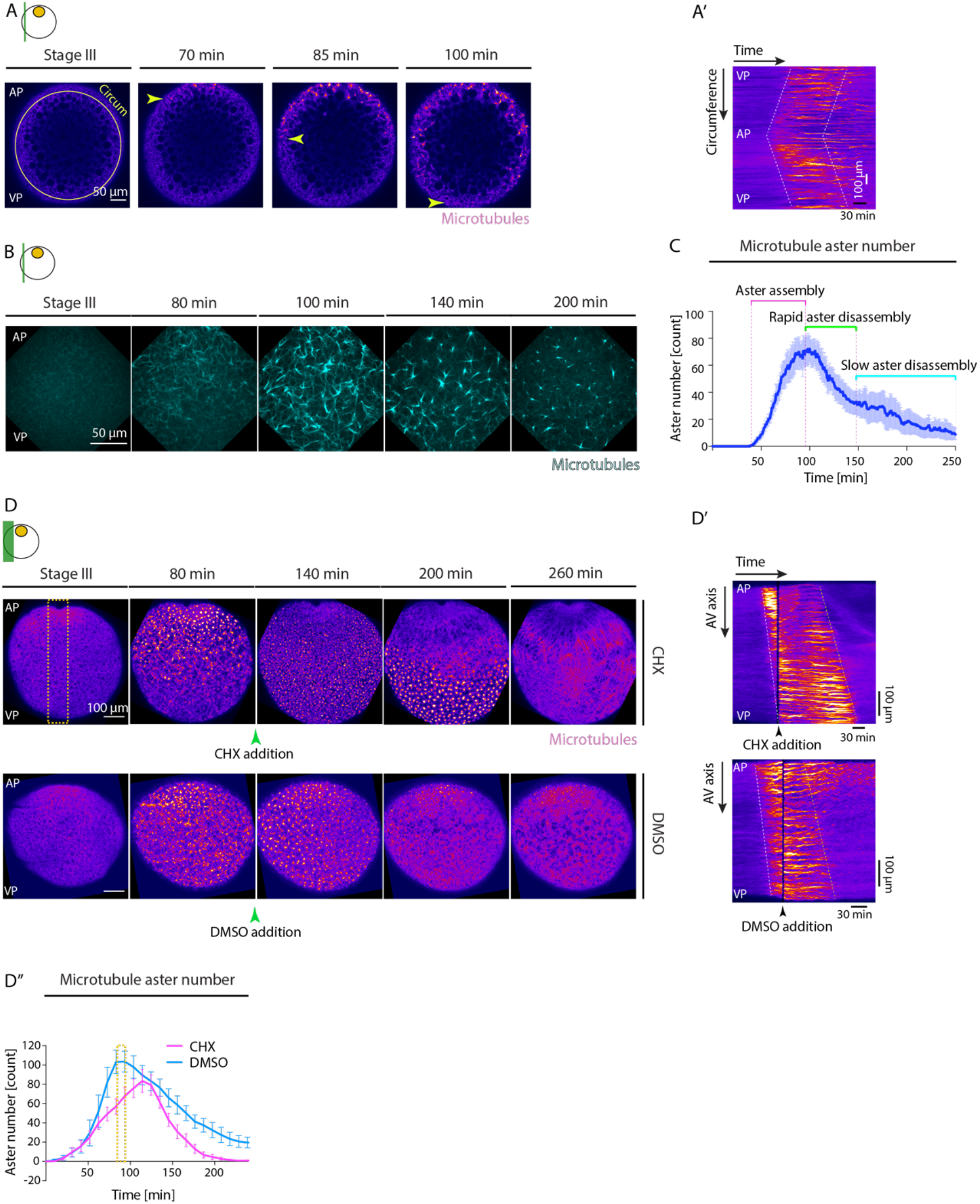
Microtubule network undergoes pronounced changes during oocyte maturation. (**A**) Fluorescence images of stage-III Tg(*Xla.Eef1a1:dclk2a-GFP*) oocytes labeling microtubules before (stage-III) and 70, 85 and 100 min after maturation onset. Arrowheads mark the propagating front of the microtubule aster formation wave. The circumferential (Circum) line in the first panel indicates the region used for acquiring the kymograph in (A’). AP, animal pole; VP, vegetal pole. (**A’**) Kymograph acquired along the circumference of the oocyte shown in (A) as a function of time. Dashed lines mark the leading and trailing edges of the microtubule aster formation wave. (**B**) Maximum fluorescence intensity projection of high resolution images of stage-III Tg(*Xla.Eef1a1:dclk2a-GFP*) oocytes labeling microtubules during consecutive stages before (stage-III) and 80, 100, 140 and 200 min after maturation onset. (**C**) Microtubule aster number as a function of time. Aster assembly during the 2nd hour after maturation onset is followed by initially rapid and consecutively slow disassembly phases (N=1 experiment, n=5 oocytes). (**D**) Maximum fluorescence intensity projection of stage-III Tg(*Xla.Eef1a1:dclk2a-GFP*) oocytes labeling microtubules exposed to Cycloheximide (CHX, inhibiting CyclinB synthesis, top row) or DMSO (control, bottom row) before and 80, 140, 200 and 260 min after maturation onset. Arrowheads mark the time point of oocyte exposure to CHX/DMSO. The dashed box indicates the animal-vegetal (AV) axis used for acquiring the kymographs in (D’). (**D’**) Kymographs acquired along the AV axis of oocytes exposed to CHX (top) or DMSO (control, bottom) as a function of time. Arrowheads mark the time point of oocyte exposure to CHX/DMSO. (**D’’**) Microtubule aster number as a function of time for oocytes exposed to CHX (magenta, N=2, n=9) and DMSO (cyan, N=2, n=8). The yellow box indicates the time window of oocyte exposure to DMSO or CHX in the respective experiments. Schematics in each panel demarcate the imaging plane used for obtaining the images in that panel. Error bars, SEM.

We next asked what signals might trigger this microtubule transformation wave. Given that the microtubule transformation wave was initiated upon GV breakdown and that the cell cycle regulator CyclinB/Cdk1 complex, stored within GV (Fig. 2E), has previously been found to trigger microtubule reorganization (Belmont et al., 1990; Ishihara et al., 2014; Verde, 1991; Verde et al., 1992), we hypothesized that the release of CyclinB/Cdk1 at the animal pole upon GV breakdown and its diffusion towards the vegetal pole of the oocyte might be involved in this process. To test this possibility, we exposed stage-IV oocytes undergoing maturation to 700 μM Cycloheximide inhibiting the synthesis of CyclinB or to 250 μM of the Cdk1 inhibitor Dinaciclib. Strikingly, microtubule asters prematurely disappeared in oocytes exposed to Cycloheximide or Dinaciclib, leading to a more homogeneous distribution of microtubules reminiscent of the situation in immature oocytes before GV breakdown (Fig. 3D-D” and S4B and video S6). This suggests that CyclinB/Cdk1 activation and release upon GV breakdown drives the observed microtubule transformation wave.

### Partial microtubule depolymerization drives microtubule aster formation during oocyte maturation

But how is the CyclinB/Cdk1 complex driving microtubule aster formation? The formation of aster-like acentrosomal microtubule structures in *Xenopus* egg extracts has previously been shown to rely on the activity of microtubule motor dynein, clustering microtubule minus ends to aster centers (Foster et al., 2015). Hence, we examined whether dynein motors might also function in driving the microtubule network reorganization observed during zebrafish oocyte maturation. To this end, we exposed stage-III oocytes to 75 μM of the dynein inhibitor Ciliobrevin D (Cheng and Ferrell, 2019). Microtubule asters of Ciliobrevin D-treated oocytes failed to fully contract and either generated abnormal asters or structures remaining connected across large distances within the oocyte (Fig. S4C-C’), suggesting that dynein motor activity is needed for proper microtubule aster formation.

To further determine whether changes in dynein motor activity, due to the release of CyclinB/Cdk1 upon GV breakdown, might be responsible for microtubule aster formation, we first estimated the motor force regime underlying this microtubule network contraction/aster formation dynamics. To this end, we measured the contraction length scales by identifying the domain that gives rise to each microtubule cluster detectable at the final time point of fusion/contraction, and determined the size of the first and second biggest domains (*ξ_1_* and *ξ_2_*), as used in percolation theory to describe the mode of network contraction (Alvarado et al., 2013). In the local contraction regime, *ξ_1_* and *ξ_2_* are of similar magnitude, while for large-scale contractions, *ξ_1_* becomes close to the system size at the expense of *ξ_2_* shrinking in size. To explore the *ξ_1_* versus *ξ_2_* space more systematically, we performed a network contraction analysis in oocytes exposed to DMSO (control), 75 μM Ciliobrevin D to inhibit dynein motor activity or to 12.5 μM and 50 μM Taxol stabilising microtubules to various extents. This analysis indicated that control as well as Taxol-treated oocytes exhibit local contractions (*ξ_1_* ≈ *ξ_2_*), with increasing Taxol concentrations resulting in gradually larger contractile domains and consequently bigger asters (Fig. 4A-A’ and S4D and video S7). In contrast, 40% of maturing oocytes exposed to 75 μM Ciliobrevin D exhibited large-scale network contractions (*ξ_1_* > *ξ_2_*), where the size of the biggest cluster reaches close to the system size (*ξ_1_* ≈ 200 μm). In line with predictions from active gel contractility (Alvarado et al., 2013), such behavior suggests that the dynein motor activity levels is high within maturing *wild type* oocytes, ensuring local network contractions and ultimately the formation of small-sized asters.

Given that microtubule aster formation, in addition to activating dynein, can also be driven by increasing the ratio of microtubule motors to microtubule number (Roostalu et al., 2018), we asked whether a reduction in microtubule number upon GV breakdown, and thus an increase in the ratio of microtubule motors to microtubule number, might be responsible for the observed aster formation of the microtubule network. To this end, we analyzed whether the total amount of polymerized microtubules changes upon GV breakdown by monitoring intensity changes in the vicinity of the GV in oocytes of Tg(*XlEef1a1:dclk2-GFP*) fish labeling microtubules (Fig. 4B and S4E-E’ and video S8). This analysis showed that the total amount of polymerized microtubules decreased up to around 30% of its initial levels just as the first microtubule asters appeared in the ooplasm (Fig. 4B’). To determine whether such depolymerization of microtubules would indeed be sufficient to trigger this transformation, we treated immature stage-III oocytes, still displaying uniform microtubule distribution (Fig. 3A), to 300 μM of the microtubule depolymerizing drug Colchicine or DMSO as control (Fig. 4C and video S8) and analyzed resultant changes in microtubule network organization. Remarkably, partial depolymerization of microtubules in Colchicine-treated stage-III oocytes led to the premature formation of numerous microtubule asters displaying extensive fusion and splitting dynamics, while the microtubule network of DMSO-treated control oocytes remained unchanged (Fig. 4C-C’). Interestingly, the network contraction analysis of the microtubule asters forming in Colchicine-treated immature oocytes revealed small contraction length scales, indicative of the high dynein activity present already within the immature oocytes before GV breakdown (Fig. 4A-A’). This suggests that the partial depolymerization of microtubules, rather than increased dynein motor activity, upon GV breakdown is responsible for the microtubule network transformations observed during oocyte maturation.

**Figure 4.**
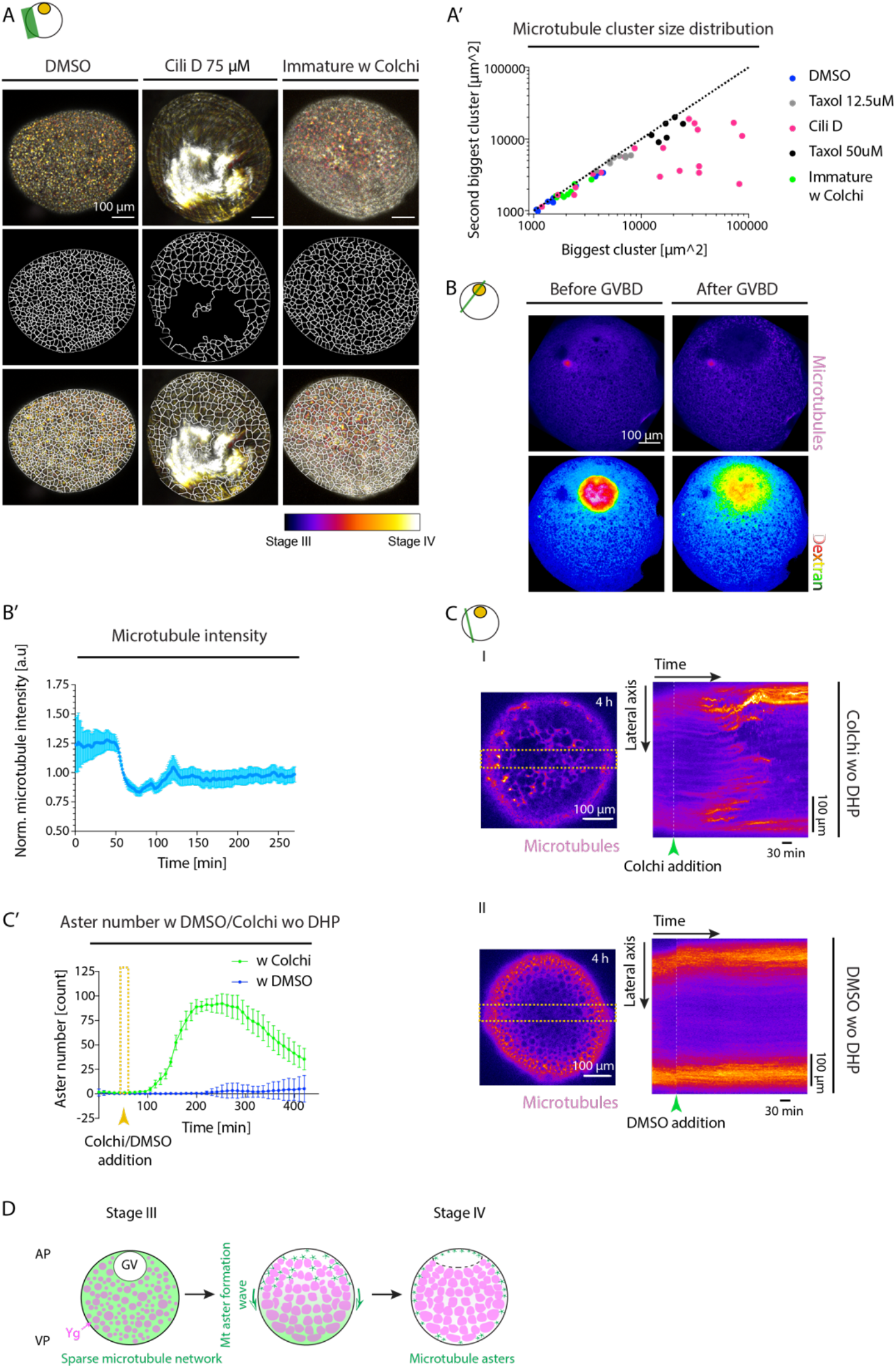
Microtubule network reorganization relies on dynein motor activity and microtubule depolymerization. (**A**) First row: Temporal maximum projection of Tg(*Xla.Eef1a1:dclk2a-GFP*) oocytes labeling microtubules and exposed to DMSO with DHP (first column), Colchicine (Colchi) without DHP (immature oocyte, second column) and 75 μM Ciliobrevin D with DHP (Cili D, third column). Second row: Voronoi triangulation of the microtubule networks shown in the first row, delimited by black lines. Third row: The overlay of the microtubule networks and their corresponding Voronoi triangulation. (**A’**) Size of the first and second biggest microtubule clusters for oocytes exposed to DMSO with DHP (blue, N=2 experiments, n=9 oocytes), 75 μM Cili D with DHP (magenta, N=2, n=19), 12.5 μM Taxol with DHP (gray, N=2, n=8), 50 μM Taxol with DHP (black, N=1, n=6) or Colchi without DHP (green, immature oocytes, N=2, n=8) obtained from images in (A) and (Fig. S4D). Black dashed line corresponds to the size of the first and second clusters being equal. (**B**) Fluorescence images of stage-III Tg(*Xla.Eef1a1:dclk2a-GFP*) oocytes labeling microtubules (first row) injected with Dextran Alexa 647 marking nucleoplasm within the germinal vesicle (GV) (second row) before and after GV breakdown (GVBD). (**B’**) Normalized microtubule intensity measured at the blastodisc region in the vicinity of the GV over time as shown in (Fig. S4E-E’), (N=2, n=9). (**C**) Fluorescent image of stage-III Tg(*Xla.Eef1a1:dclk2a-GFP*) oocytes labeling microtubules exposed to Colchi (I-left) or DMSO (control, II-left) for 4 h in the absence of DHP (immature oocyte). The orange dashed boxes denote the lateral regions used for obtaining the kymographs shown on the right. I-right and II-right: Kymographs of microtubule intensity along the lateral axis of the oocytes shown on the left as a function of time. The green arrowheads denote the time point of oocyte exposure to Colchi/DMSO. (**C’**) Microtubule aster number as a function of time for oocytes exposed to Colchi (green, N=2, n=11) or DMSO (blue, N=2, n=10) without DHP (immature oocytes). The arrowhead denotes the time point of oocyte exposure to Colchi/DMSO. (**D**) Schematic summarizing the microtubule network transformation taking place during oocyte maturation. A sparse microtubule network encompassing the ooplasm of the immature oocytes reorganizes into numerous microtubule asters in a wave-like fashion from the animal pole (AP) to the vegetal pole (VP) of the oocyte. This network transformation relies on the partial depolymerization of the microtubule network initiated by GVBD at the AP of the oocyte. Schematics in each panel demarcate the imaging plane used for obtaining the images in that panel. Error bars, SEM.

Collectively, these findings put forward a model where the microtubule network in early stage-III oocytes is in a ‘jammed’ configuration that cannot reorganize despite its high motor activity. The partial disassembly of this network upon GV breakdown will, in turn, ‘unjam’ the network by increasing the ratio of microtubule motors to microtubules, eventually leading to microtubule aster formation (Fig. 4D).

### Microtubule asters ensure proper chorion elevation upon activation

To determine whether and how the transformation of the microtubule network into acentrosomal asters is linked to its apparent requirement for chorion elevation, as suggested by our microtubule interference experiments (Fig. S2B-B’), we analyzed the spatiotemporal dynamics of microtubule aster formation relative to the reorganization of Ygs, Cgs and ooplasm. This analysis revealed that microtubule asters, while traveling as a wave from the animal to the vegetal pole of the oocyte, exhibited a radially outward-directed flow from center to the surface of the oocyte (Fig. 5A), leading to microtubule aster accumulation in cortical regions of the oocyte (Fig. 5A and video S5). Notably, the outward flow of microtubule asters was also observed in immature oocytes treated with Colchicine to trigger premature microtubule depolymerization-mediated aster formation (Fig. 4C), suggesting that this outward flow is due to some inherent radial polarity of the oocyte. This outward movement of microtubule asters might be triggered by higher levels of dynein at the cortex preferentially pulling microtubules towards the oocyte surface (Kotak et al., 2012). In line with this assumption, in oocytes exposed to Ciliobrevin D with abnormal aster formation, the microtubule asters failed to completely move towards the cortex and partially remained within the oocyte center (Fig. S4F-F’ and video S10).

**Figure 5.**
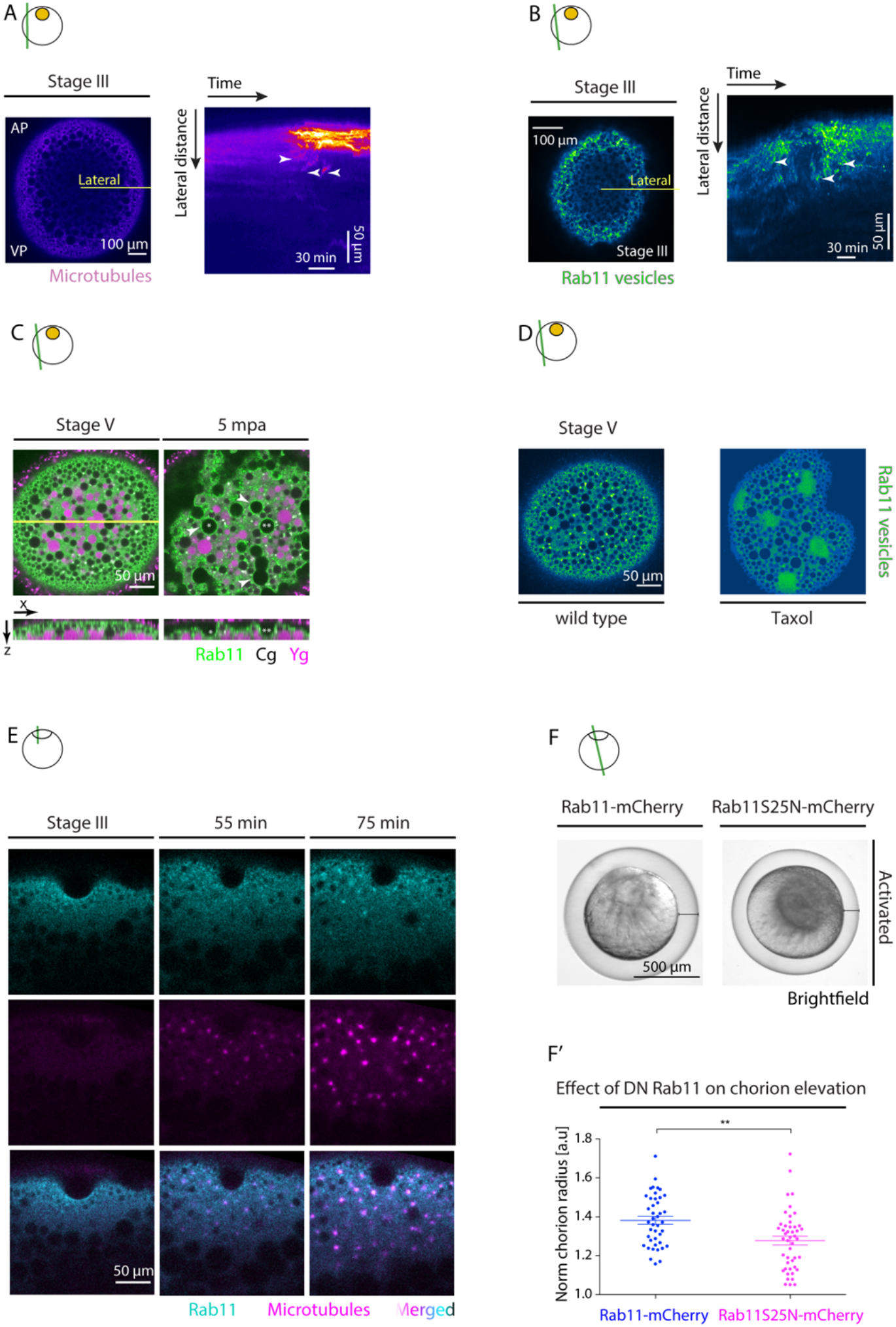
Rab11 vesicles move towards the cortex via microtubule asters. (**A**) Left: Fluorescence images of stage-III Tg(*Xla.Eef1a1:dclk2a-GFP*) oocytes labeling microtubules. AP, animal pole; VP, vegetal pole. Same oocyte as in (Fig. 3A). The lateral line indicates the region used for acquiring the kymograph in right. Right: Kymograph of microtubule intensity along the lateral axis of the oocyte shown on the left as a function of time. Arrowheads indicate exemplary outward flowing microtubule asters. (**B**) Left: Fluorescence image of stage-III Tg(*actb2:Rab11a-NeonGreen*) oocytes marking Rab11^+^ vesicles. The lateral line indicates the region used for acquiring the kymograph on the right. Right: Kymograph acquired along the lateral distance of the oocyte shown on the left as a function of time. Arrowheads point at exemplary Rab11^+^ vesicles moving towards the cortex. (**C**) Fluorescence images of Tg(*actb2:Rab11a-NeonGreen*) oocytes marking Rab11^+^ vesicles (green) and exposed to Lysotracker to label yolk granules (Yg, magenta) at stage-V (mature oocyte, left) and 5 minutes after activation (mpa) with E3 medium (right). Cortical granules (Cg, black) are identified by their exclusion of Lysotracker and the ooplasmic signal. The yellow line indicates the region used for displaying the orthogonal view (bottom images). Asterisks mark exemplary Cg undergoing exocytosis and arrowheads denote the localization of Rab11 on Cg prior to their exocytosis. (**D**) Fluorescence images of stage-V Tg(*actb2:Rab11a-NeonGreen) wild type* (WT) oocytes (top) or oocytes exposed to 50 μM Taxol (bottom). (**E**) Fluorescence images of stage-III Tg(*actb2:Rab11a-NeonGreen*) oocytes marking Rab11^+^ vesicles (green) injected with 450 pg of *DCLK-mKO2* mRNA to label microtubules (magenta) before (stage-III), and 55 and 75 min after maturation onset. (**F**) Brightfield images of oocytes injected with 350 pg of *Rab11-mcherry* (left) or *Rab11S25N-mCherry* (right; DN, dominant negative) mRNA, induced to undergo oocyte maturation for 4.5 h and activated consequently by exposure to E3 medium for 30 min. Black lines demarcate the distance between the egg and its overlaying chorion. (**F’**) Chorion elevation, normalized to the oocyte diameter, of oocytes injected with 350 pg of *Rab11-mcherry* (blue, control, N=3 experiments, n= 42 oocytes) or *Rab11S25N-mCherry* (magenta, N=3, n=46) mRNA. Schematics in each panel demarcate the imaging plane used for obtaining the images in that panel. Error bars, SEM. Mann-Whitney test, **p = 0.001.

Given the similarity in radially outward-directed movement of both microtubule asters and Cgs during oocyte maturation (Fig. 1E and 5A), we asked whether microtubules might drag Cgs towards the cortex where their exocytosis is needed for chorion elevation. However, interfering with microtubule aster formation by treating stage-III oocytes with 200 μM Colchicine did not affect Cg accumulation at the oocyte cortex (Fig. S5A-A’), suggesting that microtubules do not function in chorion elevation by transporting Cgs towards the oocyte cortex. Alternatively, microtubule aster formation and translocation towards the oocyte cortex might be required for chorion elevation by regulating Cg exocytosis. The Rab family of proteins regulates various aspects of cellular trafficking and have been implicated in Cg exocytosis in various animal species (Cheeseman et al., 2016; Sato et al., 2008; Zhen and Stenmark, 2015). In particular, Rab11 has previously been shown to localize to the surface of Cgs in both zebrafish and *C.elegans* oocytes (Kanagaraj et al., 2014; Sato et al., 2008), and to be required for synchronous secretion of Cgs upon fertilization in *C.elegans* (Sato et al., 2008). We thus hypothesized that the outward moving microtubule asters might be involved in chorion elevation by transporting Rab11-positive (Rab11^+^) vesicles towards the oocyte cortex, where Cgs reside, thereby facilitating Cg exocytosis by decorating Cgs with Rab11. To test this hypothesis, we generated Tg(*actb1:Rab11a-NeonGreen*) animals, allowing us to visualize Rab11 dynamics. Interestingly, we found that Rab11^+^ vesicles displayed outward flows during oocyte maturation, similar to microtubule asters, and eventually colocalized with Cgs at the oocyte cortex of mature oocytes (Fig. 5B-C and video S9). To determine whether microtubules are involved in this translocation of Rab11^+^ vesicles to the oocyte cortex, we analyzed whether Rab11-positive vesicle distribution changes when microtubule aster formation is altered in oocytes. To interfere with microtubule dynamics, we treated oocytes with microtubule depolymerizing (200 μM Colchicine) or stabilizing (50 μM Taxol) drugs. Taxol-treated oocytes displaying enlarged microtubule aster-like structures (Fig. S4D; Verde, 1991), exhibited large aggregates of Rab11^+^ vesicles (Fig. 5D). In contrast, oocytes exposed to Colchicine displayed strongly reduced density of Rab11^+^ vesicles at the cortex of mature oocytes (Fig. S5B-B’). To determine whether microtubule asters and Rab11-positive vesicles directly interact with each other, we simultaneously monitored Rab11^+^ vesicles and microtubules by injecting *DCLK-mKO2* mRNA, labeling microtubules, into Tg(*actb1:Rab11a-NeonGreen*) oocytes, marking Rab11^+^ vesicles (Fig. 5E and video S9). This analysis revealed that Rab11^+^ vesicles colocalized with microtubule asters as they moved towards the oocyte cortex, further supporting the notion that microtubule asters function in chorion elevation by transporting Rab11^+^ vesicles to the oocyte cortex.

Finally, to investigate whether Rab11 is indeed required for Cg exocytosis, and thus chorion elevation, we expressed a dominant-negative variant of Rab11, Rab11^S25N^ (Scheffler et al., 2021), to block Rab11 activity during oocyte maturation (Fig. 5F). Strikingly, we found that overexpression of Rab11^S25N^ led to strongly reduced chorion elevation in activated oocytes (Fig. 5F’), suggesting that Rab11 activity is required for this process. Taken together, these results indicate that microtubule asters, by moving towards the cortex upon GV breakdown, take along Rab11-positive vesicles, and that this cortical translocation of Rab11-positive vesicles is required for the decoration of Cgs at the cortex with Rab11 and, thus, Cg exocytosis.

### Yolk granule fusion triggers cytoplasmic flows transporting cortical granules towards the oocyte cortex

Questions remain as to the mechanisms underlying the relocalization of Cgs towards the oocyte cortex, since neither the inhibition of myosin II activity nor the depolymerization of microtubules affected Cg relocalization (Fig. S5A-A’ and video S11), ruling out a direct involvement of these cytoskeletal networks in this process. To identify such mechanism(s), we performed a detailed spatiotemporal analysis of Cg translocation towards the oocyte surface. This showed that in immature stage-III oocytes, Cgs were distributed in between Ygs (Fig. 1B). However, as oocyte maturation proceeded and GV breakdown occurred at the oocyte animal pole, Ygs underwent extensive fusion and compaction concomitant with Cgs translocation towards the oocyte cortex (Fig. 1B-E). Generally, the compaction of a compressible material embedded within an incompressible fluid is expected to drive outward fluid flows due to volume conservation. Therefore, we postulated that Yg fusion and compaction might result in outward ooplasmic flows, which in turn takes along Cgs, thereby translocating them to the oocyte surface. To directly test this possibility, we treated oocytes with 100 μM Ouabain, a known inhibitor of *Na*^+^/*K*^+^ ATPase pumps (Chan et al., 2019), as an increase in intracellular *K*^+^ levels has previously been suggested to induce yolk granule fusion in other teleosts (Selman et al., 2001; Wallace and Selman, 1981). In oocytes treated with Ouabain, Ygs largely failed to fuse, and, consequently, the oocyte remained opaque (Fig. 6A-A’ and S6A, video S12 and Selman et al., 2001). Moreover, Cg translocation to the cortex was strongly reduced in Ouabain-treated oocytes (Fig. 6B-B’ and video S11), suggesting that Yg fusion and the resultant ooplasmic flows are required for this process. Notably, we also observed that not only Cg translocation but also blastodisc formation was reduced in Ouabain-treated oocytes (Fig. S6B-B’ and video S10), suggesting that Yg fusion and resultant outward cytoplasmic flows are also needed for blastodisc formation. In line with such a function, we found that Yg fusion was more pronounced at the animal pole of the oocyte, where the blastodisc is forming (Fig. S6C-C’). Importantly, these different effects of Ouabain-treatment were not due to Ouabain affecting GV breakdown and/or associated actin and microtubule cytoskeletal rearrangements, as all of these processes remained unchanged in Ouabain-treated oocytes (Fig. S6D).

**Figure 6.**
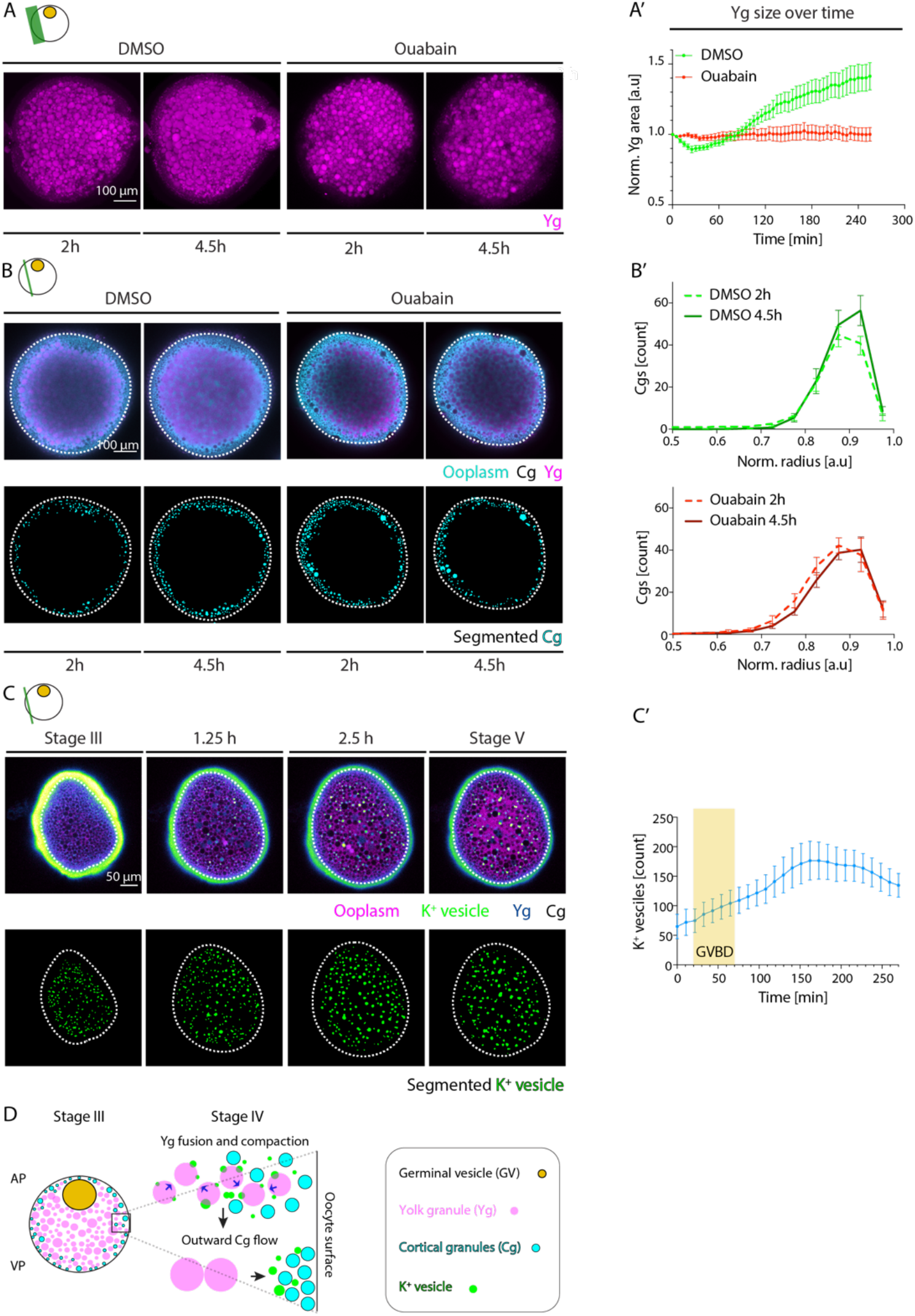
Yolk granule fusion drives cortical granules localization at the oocyte cortex. (**A**) Maximum fluorescence intensity projection of oocytes exposed to DMSO (left)/Ouabain (right) and Lysotracker for labeling yolk granules (Yg) 3h after maturation onset. (**A’**) Normalized average Yg area as a function of time during oocyte maturation for oocytes exposed to DMSO (green, N=3 experiments, n=15 oocytes) or Ouabain (red, N=3, n=14). (**B**) Fluorescence images of DMSO-treated (left)/Ouabain-treated (right) stage-III Tg(*hsp:clip170-GFP*) oocytes labeling ooplasm (cyan) and exposed to Lysotracker to mark Yg (magenta) and cortical granules (Cg, black, identified by their exclusion of both Clip-170-GFP and Lysotracker) at 2 and 4.5 h after maturation onset (top row). Images in the bottom row show segmented Cg obtained from the images in the top row. Dashed lines mark the oocyte outline. (**B’**) Average Cg density profile across the oocyte radius for DMSO- (top, green, N=3, n=11) and Ouabain- (bottom, red, N=3, n=14) treated oocytes at 2 and 4.5 h after maturation onset. Normalized (norm.) radius of 0 and 1 correspond to the oocyte center and surface, respectively. Note that changes, and not the absolute values, in Cg distribution along the oocyte radius between 2 and 4.5 h correspond to Cg movements during this time. (**C**) Top row: Fluorescence images of stage-III oocytes injected with *K*^+^ indicator (*K*^+^-Green, green), Dextran Alexa Fluor 647 to mark ooplasm (magenta) before (stage-III) and 1.25, 2.5 and 4.5 h after maturation onset. Cg (black) are identified by their exclusion of both Dextran and *K*^+^-Green. The vesicles enriched with *K*^+^ fused with each other and Yg, thereby increasing their internal *K*^+^ concentration and hence becoming dark blue (in Green-Fire-Blue Look Up Table). Bottom row: Segmented *K*^+^ vesicles identified from the images in the top row. Dashed lines mark the oocyte outline. (**C’**) Number of *K*^+^ vesicles in superficial stacks of the oocyte as in (C) as a function of time (N=2, n=8). Yellow box indicates the period during which GVBD takes place. (**D**) Schematic summarizing the role of Yg fusion in ooplasmic reorganizations during zebrafish oocyte maturation. GV breakdown leads to Yg fusion by triggering a *Na*^+^/*K*^+^ ATPase-dependent increase in the concentration of *K*^+^ within Ygs. The fusion and compaction of Ygs to the oocyte center, in turn, squeezes the ooplasm outward towards the oocyte surface. These ooplasmic flows, in turn, carry along Cgs, thereby moving them closer to the cortex. Schematics in each panel demarcate the imaging plane used for obtaining the images in that panel. Error bars, SEM.

Taken together, these results suggest that Yg fusion and its associated radially outward-directed ooplasmic flows drive translocation of Cg to the oocyte cortex and contribute to blastodisc formation at the oocyte animal pole.

Finally, we asked what signals might trigger Yg fusion. Given that an increase in intracellular *K*^+^ levels has previously been proposed to trigger YG fusion in Sea Bass oocytes (Selman et al., 2001), we monitored dynamic changes in intracellular *K*^+^ levels during oocyte maturation by injecting a *K*^+^ indicator (*K*^+^-Green) into the immature stage-III oocyte. Strikingly, we found *K*^+^ to become enriched in small vesicles, which increased in number upon GV breakdown and underwent extensive fusion with each other and Ygs, ultimately increasing *K*^+^ levels inside Ygs undergoing fusion (Fig. 6C-C’). In contrast, no such spatiotemporal changes in *K*^+^ levels upon GV breakdown were detected in Ouabain-treated oocytes defective in Yg fusion (Fig. S6E-E’ and video S12). Given that Ouabain blocks *Na*^+^/*K*^+^ ATPase pumps, this suggests that GV breakdown leads to Yg fusion by triggering a *Na*^+^/*K*^+^ ATPase-dependent increase in the concentration of *K*^+^ within Ygs. This Yg fusion and compaction to the oocyte center, in turn, triggers radially outward-directed ooplasmic flows, which carry along and position Cgs at the oocyte cortex (Fig. 6D).

## Discussion

Our study provides novel insight into both the processes underlying oocyte maturation and the general mechanisms by which cytoplasmic reorganization is achieved within cells. Cortical granules are Golgi-derived secretory vesicles, which localize to the cortex of the mature oocytes to undergo exocytosis upon fertilization and prevent polyspermy (Liu, 2011). Small Rab GTPase family of proteins have been found to associate with cortical granules and regulate their transport and exocytosis in oocytes of diverse organisms (Cheeseman et al., 2016; Rojas et al., 2021; Sato et al., 2008). In mouse oocytes, for instance, positioning of cortical granules to the oocyte cortex has been proposed to rely on both myosin Va motors localizing to these granules and inducing their movement on an intrinsically polarized bulk actin network, as well as “hitchhiking” on outward moving Rab11a-vesicles (Cheeseman et al., 2016). Our findings that yolk granule fusion and compaction to the oocyte center drive cortical granule movements towards the oocyte circumference, and that microtubule aster formation and translocation to the cortex lead to the concomitant accumulation of Rab11 positive vesicle at the circumference, identifies a yet unknown mechanism of cortical granule translocation and exocytosis during oocyte maturation. Why the mechanisms regulating cortical granule translocation and exocytosis differ between mice and zebrafish is still unknown, but likely the presence of yolk granules in zebrafish but not mouse oocytes has led to different functional adaptations of cytoplasmic components.

Beyond the specific regulation of cortical granule localization within the maturing oocyte, our findings also shed light on the general mechanisms underlying cytoplasmic reorganization and the role of the cell cytoskeleton therein. Cytoskeletal networks, such as the actin and microtubule cytoskeleton, have been implicated as the main driving force underlying cytoplasmic organization in oocytes and embryos. In particular, myosin II-mediated actin network flows dragging the adjacent cytoplasm have been shown to generate cytoplasmic streaming in early *Drosophila* and zebrafish embryos (Deneke et al., 2019; Shamipour et al., 2019), and the motion of kinesin I motors along microtubule arrays that are anchored to the cortex to drive cytoplasmic flows in *Drosophila* and *C. elegans* oocytes by exerting viscous drag forces to the surrounding ooplasm (Kimura et al., 2017; Monteith et al., 2016). In addition, the assembly and interaction of actin comets and centrosomal microtubule asters with organelles such as yolk granules, the nucleus or the meiotic spindle can position those organelles to specific locations within oocytes and embryos (Li et al., 2008; Shamipour et al., 2019; Theriot et al., 1992). Our findings that the partial disassembly of a pre-stressed microtubule network results in its transformation into acentrosomal microtubule asters, which move towards the oocyte cortex in a dynein-dependent manner, thereby carrying along and enriching Rab11^+^ vesicles at the oocyte surface, suggest a novel mechanism by which the microtubule network can affect vesicle localization within cells. Moreover, our observation that such transformation of the microtubule network is driven by the cell cycle regulators Cdk1/CyclinB partially disassembling the microtubule network, mechanistically links cell cycle progression to cytoskeletal reorganization in cells.

One of the key findings of our study is that the regulated fusion of yolk granules can trigger outward cytoplasmic flows, which in turn transport cortical granules towards the oocyte circumference. This suggests that beyond ATP/GTP hydrolysis-dependent actin/microtubule polymerization and the activity of their associated motor proteins, large-scale cytoplasmic flows and reorganization can also be achieved by coordinated changes in the size and shape of organelles. Interestingly, these are processes that - differently from the aforementioned cytoskeletal rearrangements - do not directly depend on the activity of motor proteins and polymerases, but instead are regulated by changes in the propensity of organelles to deform and/or undergo fusion and fission. Our observation that yolk granule fusion is mediated by a rise in the level of K^+^ within yolk granules, which depends on the activity of Na^+^/K^+^ ATPase pumps and is triggered by GV breakdown, points at intracellular ion concentration as a critical regulator of organelle fusion and resultant cytoplasmic reorganization. Notably, an increase of intracellular K^+^ within the oocyte coinciding with the fusion of yolk granules has previously been observed in oocytes of various teleosts, suggesting that K^+^-driven yolk granule fusion might represent an evolutionary conserved mechanism in yolk granule-containing oocytes. How GV breakdown leads to an increase in K^+^ levels within yolk granules, and how such increase promotes yolk granule fusion is not yet understood, but it is conceivable that electrostatic interactions between the plasma membrane surrounding yolk granules and K^+^ molecules promote the capacity of their plasma membrane to undergo fusion.

Cytoplasmic reorganization relies on the inherent self-organizing properties of the cytoplasm and a combination of pre-patterning of the cell and external signals modulating this self-organizing activity. While the external signals and pre-patterning can vary depending on the specific organismal context, cytoplasmic self-organization represents a generic propensity of the cytoplasm that emerges as a result of a certain composition or state of the cytoplasm. Our finding of yolk granule fusion eliciting large-scale cytoplasmic streaming reveals a novel feature of such inherent self-organizing capacity of the cytoplasm that plays an important role in reorganizing the cytoplasm during zebrafish oocyte maturation. Given that yolk granules constitute a major cytoplasmic component in oocytes from birds, reptiles, worms, and fish, yolk granule fusion might represent a common principle of regulating cytoplasmic streaming during oogenesis.

## Supplementary Information

**Figure S1.**
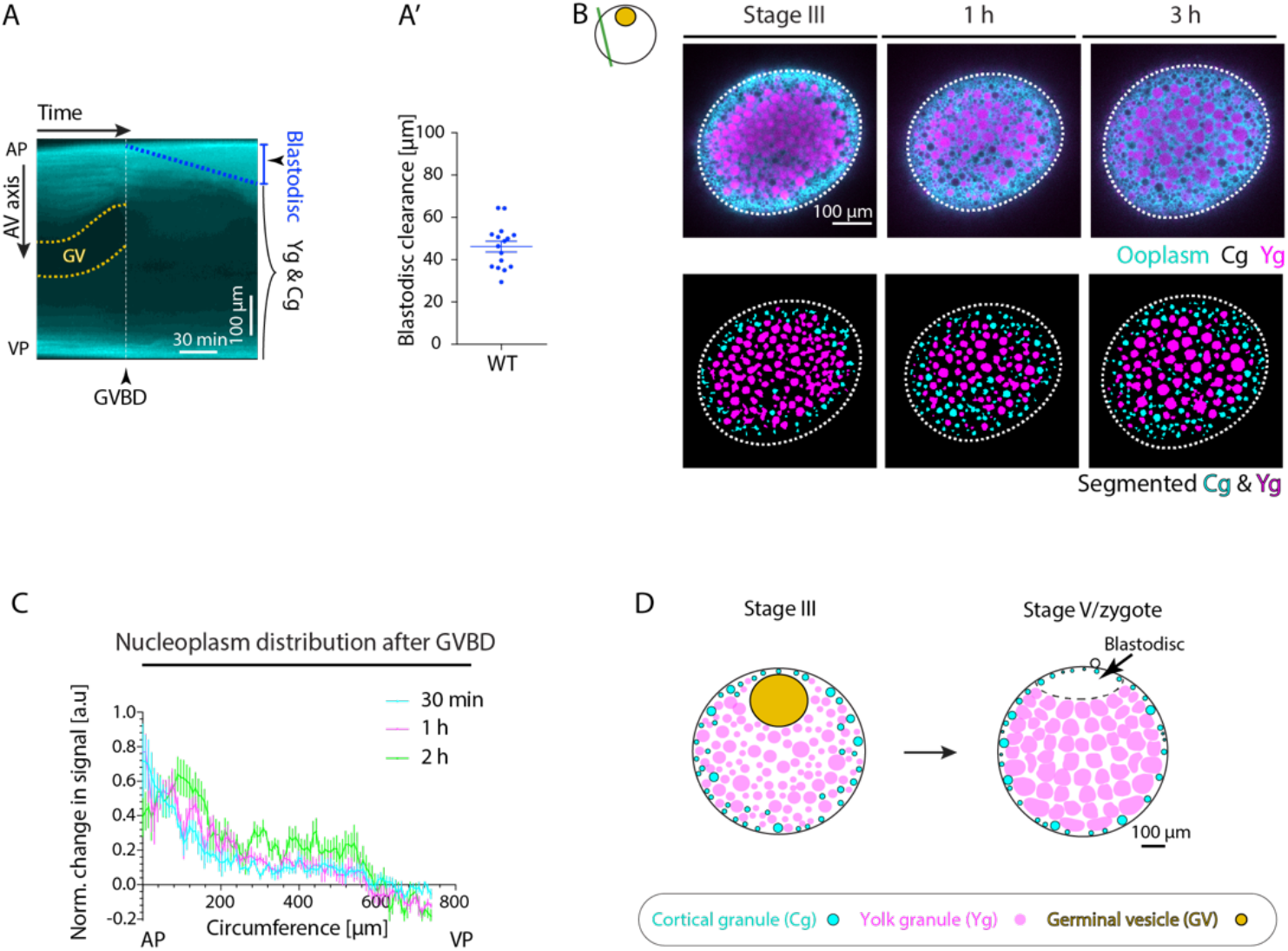
(**A**) Kymograph acquired along the animal-vegetal (AV) axis of the oocyte shown in Fig. 1A as a function of time. The dashed white line marks the time point of germinal vesicle breakdown (GVBD). The yellow dashed lines outline the germinal vesicle (GV) contour. The blue dashed zone demarcates the blastodisc region. Yg, yolk granules; Cg, cortical granules. (**A’**) Blastodisc clearance, measured as the height of blastodisc at the end of the maturation process as shown in Fig. S1A, for *wild type* (WT) oocytes (N=2 experiments, n=16 oocytes). (**B**) Fluorescence images of stage-III oocytes expressing Tg(*hsp:clip170-GFP*) to mark ooplasm (cyan) and exposed to Lysotracker to distinguish Yg (magenta) from Cg (black, identified by their exclusion of both Clip-170-GFP and Lysotracker) before (stage-III) and 1 and 3 h after maturation onset (top row). Images in the bottom row show segmented Cg obtained from the images in the top row. Dashed lines demarcate the oocyte outline. (**C**) Normalized change in Dextran Alexa 647 signal injected to label GV nucleoplasm, relative to its distribution at the time point before GVBD, at 30 min (cyan), 1 h (magenta) and 2 h (green) after GVBD plotted as a function of the oocyte circumference (N=1, n=6). Circumference of 0 and 1 correspond to the oocyte animal pole (AP) and vegetal pole (VP), respectively. (**D**) Schematic summarizing the ooplasmic reorganizations occurring during zebrafish oocyte maturation. At the onset of maturation, the GV (yellow) breaks down, triggering blastodisc formation at the AP of the oocyte. Concomitantly, Yg (magenta) fuse and compact towards the oocyte center, while Cg (cyan) translocate to the oocyte surface. Schematics in each panel demarcate the imaging plane used for obtaining the images in that panel. Error bars, SEM.

**Figure S2.**
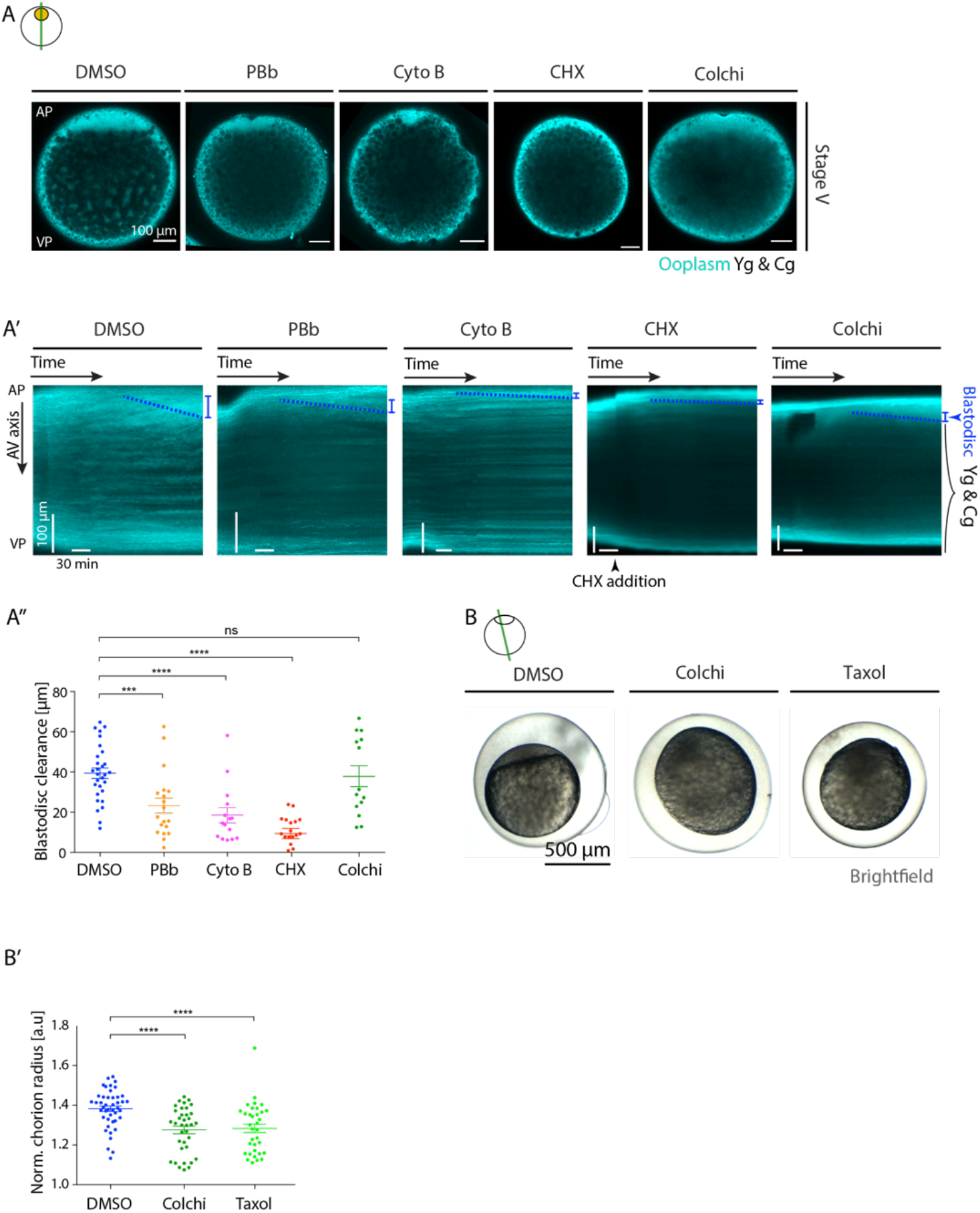
(**A**) Fluorescence images of stage-V Tg(*hsp:clip170-GFP*) oocytes labeling ooplasm (cyan) exposed to DMSO (control), 100 μM para-Nitroblebbistatin (PBb, inhibiting myosin II activity), 30 μg/ml Cytochalasin B (Cyto B, blocking actin polymerization), 700 μM Cycloheximide (CHX, blocking CyclinB synthesis, added at 45 min after maturation onset) or 200 μM Colchicine (Colchi, inhibiting microtubule polymerization). AP, animal pole; VP, vegetal pole. Yg, yolk granules; Cg, cortical granules. (**A’**) Kymographs acquired along the animal-vegetal (AV) axis of the oocytes shown in (A) as a function of time. The blue dashed lines demarcate the blastodisc expansion, the slope of which was used to measure the blastodisc clearance in (A’’). (**A’’**) Blastodisc clearance, measured as the height of blastodisc at the end of the maturation process as shown in (A’), for oocytes exposed to DMSO (N=4 experiments, n=29 oocytes), PBb (N=3, n=19), Cyto B (N=3, n=15), Colchi (N=3, n=14) or CHX (N=2, n=18). (**B**) Brightfield images of oocytes exposed to DMSO, 250 μM Colchi or 25 μM Taxol (stabilizing microtubules) and induced to undergo oocyte maturation for 4.5 h and activated consequently by exposure to E3 medium for 30 min. (**B’**) Chorion elevation, normalized to the oocyte diameter, of oocytes exposed to DMSO (blue, control, N=3, n=44), Colchi (dark green, N=3, n=36) or Taxol (light green, N=3, n=34). Schematics in each panel demarcate the imaging plane used for obtaining the images in that panel. Error bars, SEM.

**Figure S3.**
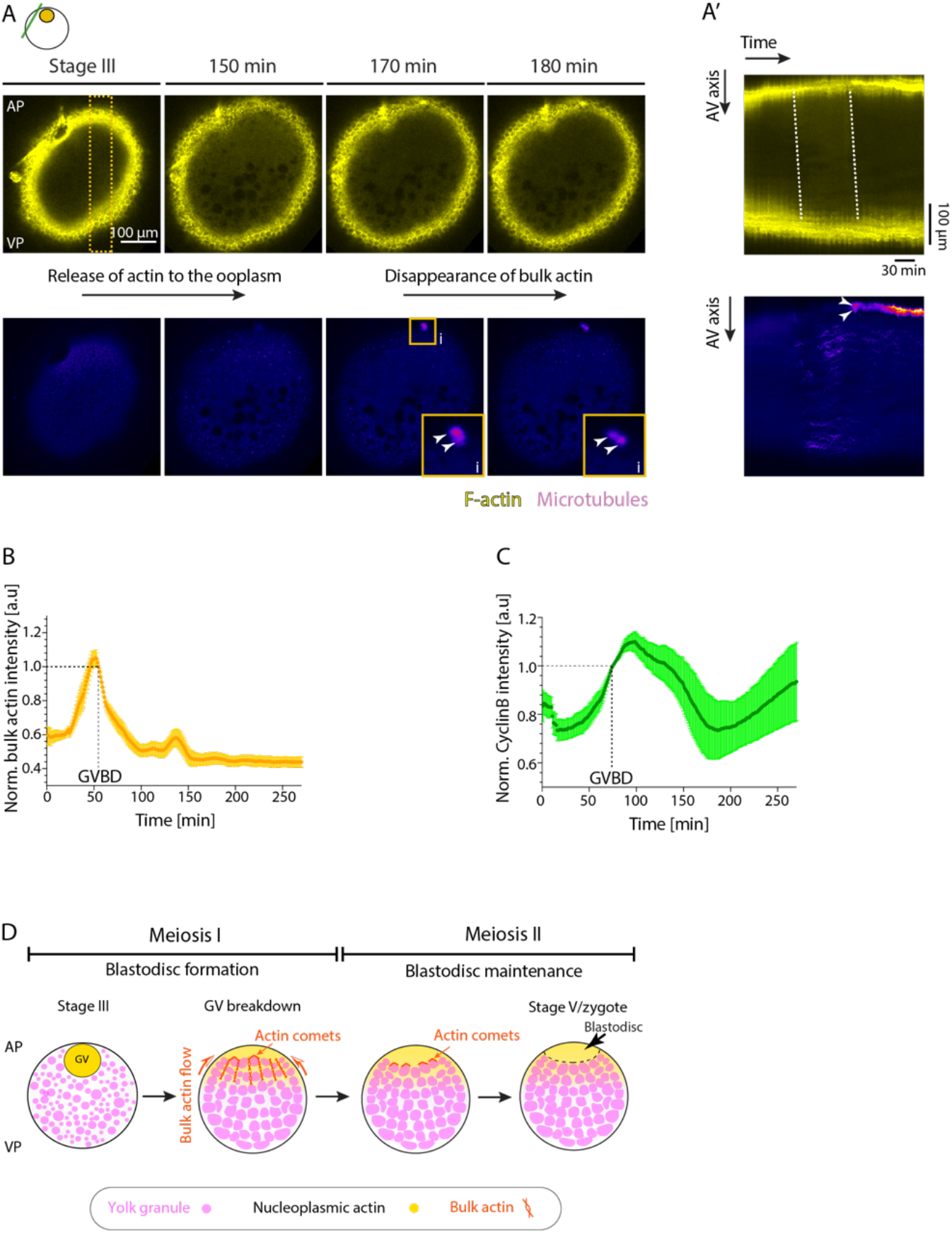
(**A**) Fluorescence images of stage-III Tg(*actb1:Utr-mCherry);(Xla.Eef1a1:dclk2a-GFP*) oocytes labeling F-actin (yellow, top row) and microtubules (purple, bottom row) before (stage-III) and 150, 170 and 180 min after maturation onset. The dashed box indicates the region used for acquiring the kymographs in (A’). AP, animal pole; VP, vegetal pole. The solid box indicates the region used for the zoomed-in view (i). White arrowheads in the insets mark the meiotic spindle formation at the end of the first meiosis. Schematic in this panel demarcates the imaging plane used for obtaining the images. (**A’**) Kymographs of F-actin (top) and microtubules (bottom) acquired along the animal-vegetal (AV) axis of the oocyte in (A) as a function of time. Arrowheads mark the first meiotic spindle. Dashed lines trace the increase of bulk actin within the ooplasm upon germinal vesicle breakdown (GVBD) and its decrease prior to completion of the first meiosis (as the first meiotic spindle formed, arrowheads). (**B**) Normalized bulk actin intensity measured at the AP of Tg(*actb1:Utr-GFP*) oocytes over time (N=2 experiments, n=9 oocytes). The dashed line denotes the timepoint of GVBD. (**C**) Normalized CyclinB intensity measured at the animal pole of the oocytes injected with *CyclinB-GFP* mRNA over time (N=2, n=7). The dashed line denotes the timepoint of GVBD. (**D**) Schematic summarizing the role of actin in ooplasmic flows and blastodisc formation. Bulk actin, initially stored within the germinal vesicle (GV), is released at the AP of the oocyte upon GVBD, thereby generating a local actin gradient. This actin gradient, in turn, directs bulk actomyosin flows towards the AP, which by dragging the ooplasm along results in blastodisc formation. In addition, the blastodisc interface is maintained during first and second meiosis by actin comet-like structures forming on the surface of yolk granules and pushing them outside of the blastodisc region. Schematic in (A) demarcates the imaging plane used for obtaining the images in this panel. Error bars, SEM.

**Figure S4.**
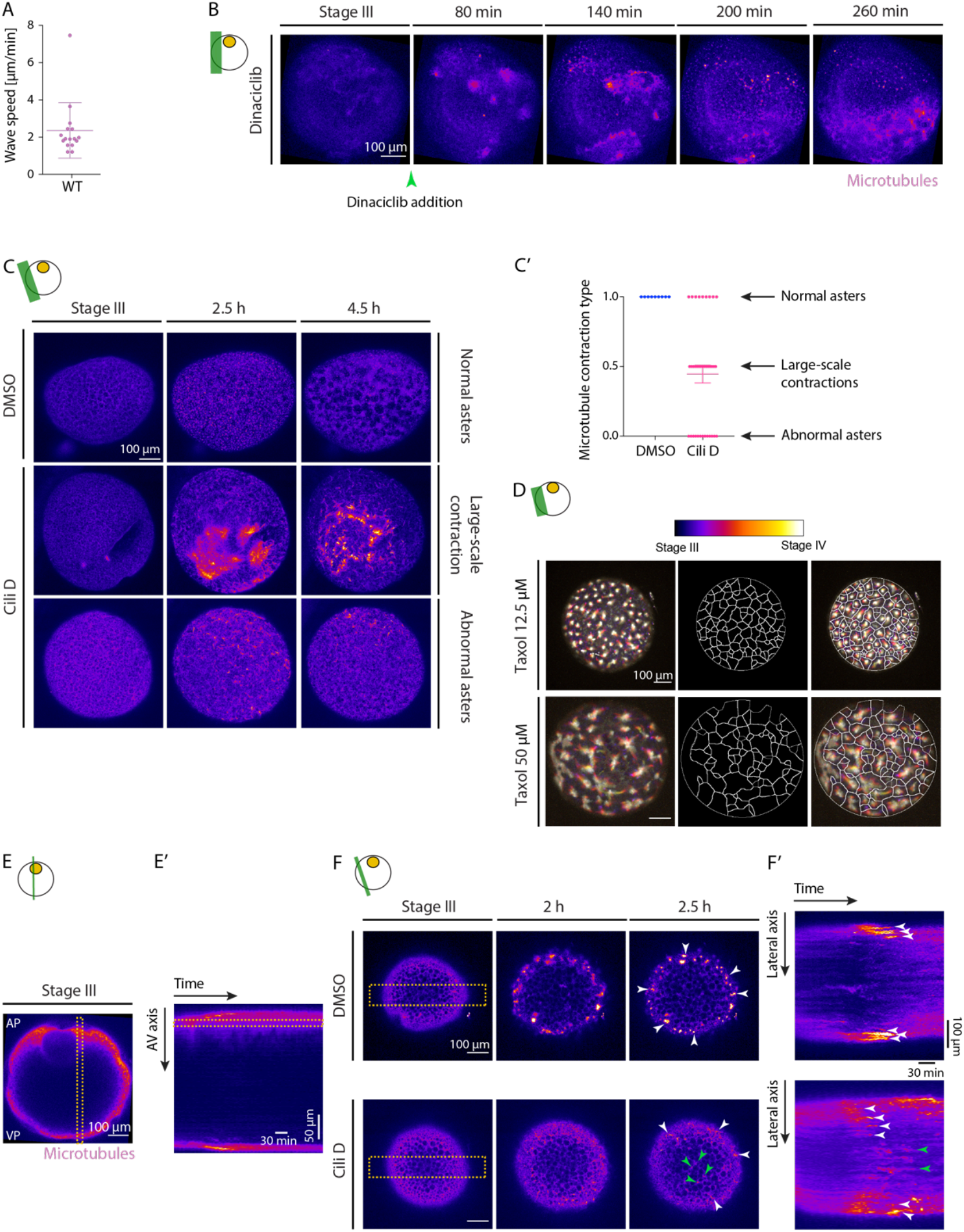
(**A**) Speed of microtubule aster formation wave along the oocyte circumference, measured from kymographs as shown in (Fig. 3A’) (N=9 experiments, n=14 oocytes). (**B**) Maximum fluorescence intensity projection of stage-III Tg(*Xla.Eef1a1:dclk2a-GFP*) oocytes labeling microtubules exposed to 250 μM Dinaciclib before (stage-III) and 80, 140, 200 and 260 min after maturation onset. Arrowhead marks the time point of oocyte exposure to Dinaciclib. (**C**) Maximum fluorescence intensity projection of stage-III Tg(*Xla.Eef1a1:dclk2a-GFP*) oocytes labeling microtubules exposed to DMSO (control, first row) or 75 μM Ciliobrevin D (Cili D, second and third rows) before (stage-III) and 2.5 and 4.5 h after maturation onset. Second row: An exemplary oocyte exposed to Cili D with abnormal microtubule aster formation. Third row: An exemplary oocyte exposed to Cili D with large-scale microtubule network contraction. (**C’**) Microtubule contraction type for DMSO-/Cili D-treated oocytes. Contraction type of 0, 0.5 and 1 correspond to abnormal asters (panel C – third row), large-scale contractions (panel C – second row) and normal asters (panel C – first row), respectively. (**D**) First row: Temporal maximum projection of Tg(*Xla.Eef1a1:dclk2a-GFP*) oocytes labeling microtubules and exposed to 12.5 μM (first column) and 50 μM (second column) Taxol in the presence of DHP. Second row: Voronoi triangulation of the microtubule networks shown in the first row, outlined by black lines. Third row: Overlay of the microtubule networks and their corresponding Voronoi triangulation. (**E**) Fluorescence image of stage-III Tg(*Xla.Eef1a1:dclk2a-GFP*) oocyte labeling microtubules. AP, animal pole; VP, vegetal pole. The dashed box indicates the region used for acquiring the kymograph on the right. (**E’**) Kymograph of microtubules acquired along the animal-vegetal (AV) axis of the oocyte shown in (E) as a function of time. The intensity profile along the dashed line was used to obtain the microtubule intensity profile shown in (Fig. 4B’). (**F**) Fluorescence images of stage-III Tg(*Xla.Eef1a1:dclk2a-GFP*) oocytes labeling microtubules exposed to DMSO (top row) or 75 μM Cili D (bottom row) before (stage-III) and 2 and 2.5 h after maturation onset. Green and white arrowheads demarcate microtubule asters below the oocyte surface (central portion of the imaging plane) and at the oocyte surface (marginal region of the imaging plane), respectively. The dashed boxes indicate the lateral regions used for acquiring the kymographs in (F’). (**F’**) Kymographs acquired along the lateral axis of oocytes exposed to DMSO (top) or Cili D (bottom) as a function of time. Green and white arrowheads demarcate microtubule asters below the oocyte surface and at the oocyte surface, respectively. Schematics in each panel demarcate the imaging plane used for obtaining the images in that panel. Error bars, SEM.

**Figure S5.**
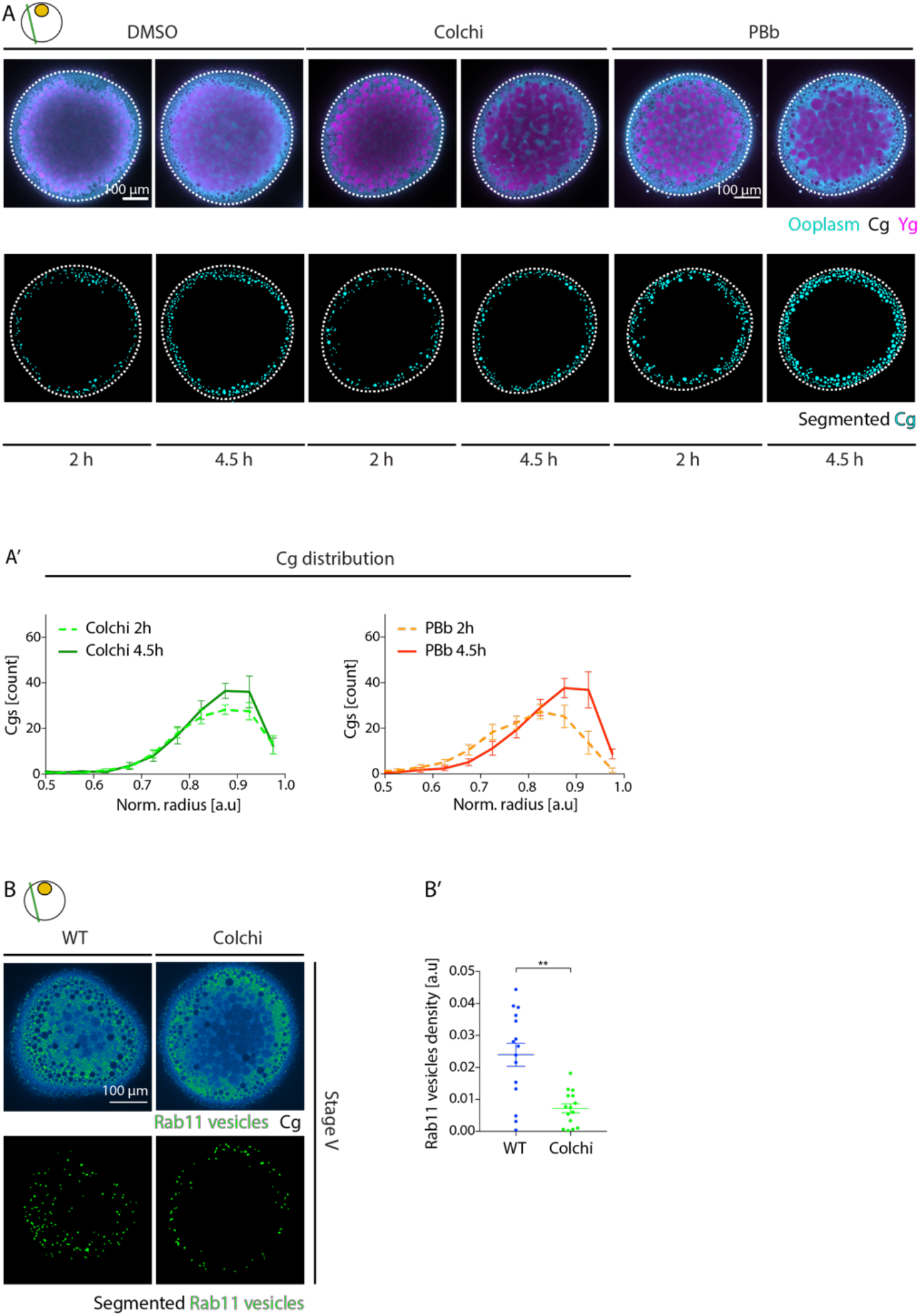
(**A**) Top row: Fluorescence images of stage-IV oocytes expressing Tg(*hsp:clip170-GFP*) to mark ooplasm (cyan) and exposed to Lysotracker to mark yolk granules (Yg, magenta) for oocytes treated with DMSO, 200 μM Colchicine (Colchi) or 100 μM para-Nitroblebbistatin (PBb). Cortical granules (Cg, black) were identified by their exclusion of both Clip-170-GFP and Lysotracker. Bottom row: Segmented Cg obtained from the images in the top row. White dashed lines mark the oocyte outline. (**A’**) Cg density profile along the oocyte radius at 2 and 4.5h after maturation onset for oocytes exposed to Colchi (left, N=3 experiments, n=15 oocytes) or PBb (right, N=2, n=12). Normalized (norm.) radius of 0 and 1 correspond to the oocyte center and surface, respectively. Note that changes, and not the absolute values, in Cg distribution along the oocyte radius between 2 and 4.5 h correspond to Cg movements during this time. (**B**) Top row: Fluorescence images of stage-V Tg(*actb2:Rab11a-NeonGreen*) oocytes. Left: *wild type* (WT), right: oocytes exposed to 200 μM Colchi. Bottom row: Segmented Rab11-positive vesicles obtained from the images in the first row. (**B’**) Density of Rab11-positive vesicles, measured from the images in (B) for WT (blue, N=2, n=15) and Colchi-treated oocytes (green, N=2, n=15). Schematics in each panel demarcate the imaging plane used for obtaining the images in that panel. Error bars, SEM. Mann-Whitney test, **p = 0.0012.

**Figure S6.**
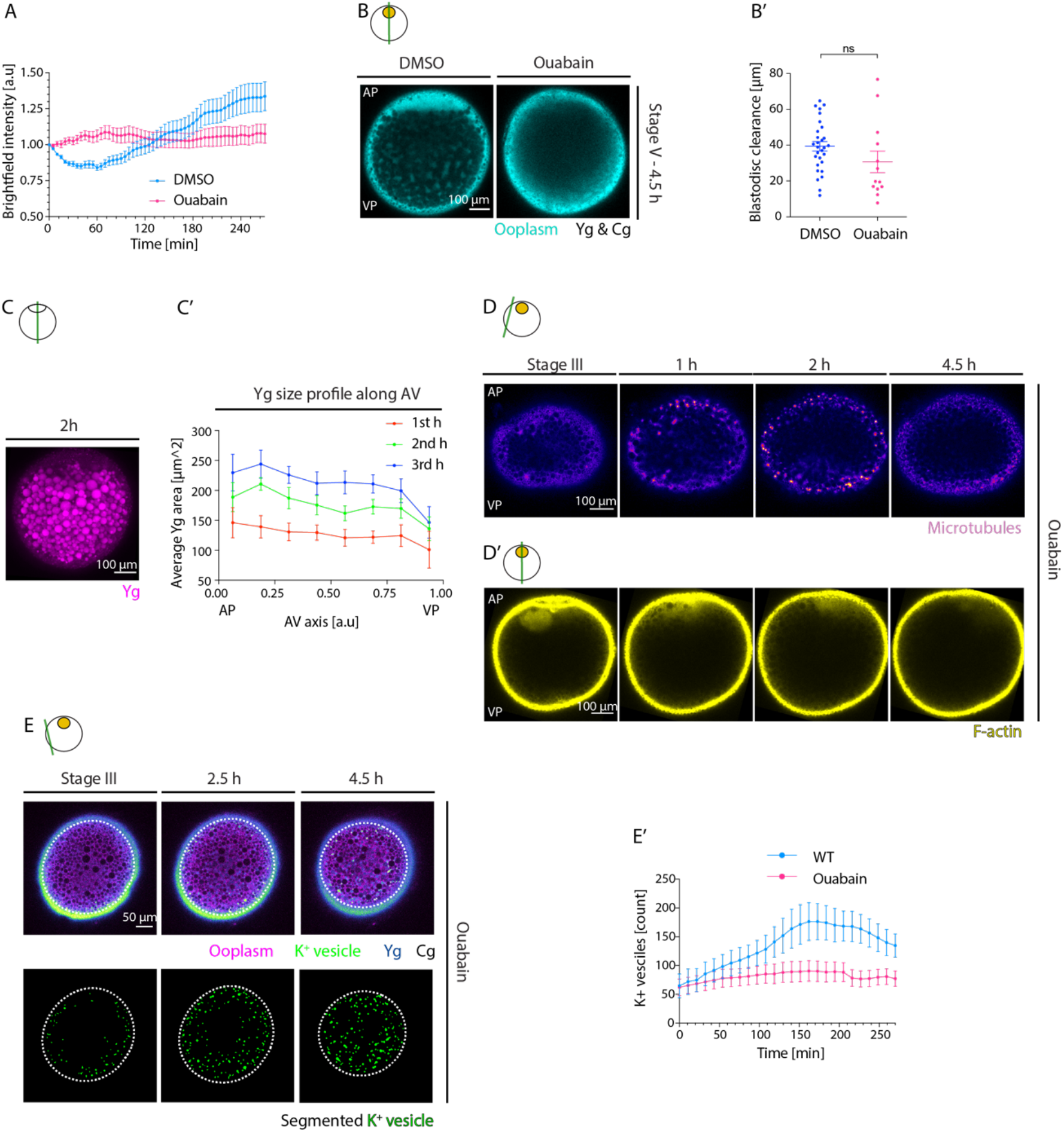
(**A**) Normalized brightfield intensity of oocytes exposed to DMSO (control, blue, N=3 experiments, n=15 oocytes) or 100 μM Ouabain (magenta, N=3, n=14) over time. (**B**) Fluorescence images of stage-V Tg(*hsp:clip170-GFP*) oocytes labeling the ooplasm and exposed to DMSO (control) or 100 μM Ouabain. AP, animal pole; VP, vegetal pole. (**B’**) Blastodisc clearance, measured as the height of blastodisc at the end of the maturation process for oocytes exposed to DMSO (control, blue, N=4, n=29) or 100 μM Ouabain (magenta, N=3, n=13). (**C**) Maximum fluorescence intensity projection of oocytes exposed to Lysotracker to mark yolk granules (Yg) 2 h after maturation onset. (**C’**) Average Yg area distribution along the oocyte animal-vegetal (AV) axis and averaged for the 1st, 2nd and the 3rd h after maturation onset (N=3, n=8). (**D**) Fluorescence images of stage-III Tg(*Xla.Eef1a1:dclk2a-GFP*) oocytes labeling microtubules exposed to 100 μM Ouabain before (stage-III) and 1, 2 and 4.5 h after maturation onset. (**D’**) Fluorescence images of stage-III Tg(*actb1:Utr-GFP*) oocytes labeling F-actin exposed to 100 μM Ouabain before (stage-III) and 1, 2 and 4.5 h after maturation onset. (**E**) Top row: Fluorescence images of stage-III oocytes injected with *K*^+^ indicator (*K*^+^-Green, green), Dextran Alexa Fluor 647 to mark ooplasm (magenta) and exposed to 100 μM Ouabain before (stage-III) and 1, 2 and 4.5 h after maturation onset. Cg (black) are identified by their exclusion of both Dextran and *K*^+^-Green. Bottom row: Segmented *K*^+^ vesicles identified from the images in the top row. Dashed lines mark the oocyte outline. (**E’**) Number of *K*^+^ vesicles in superficial stacks of the *wild type* oocytes (WT, blue, N=2, n=8, same data as in Fig. 6C’) or oocytes exposed to 100 μM Ouabain (magenta, N=2, n=12) over time. Schematics in each panel demarcate the imaging plane used for obtaining the images in that panel. Error bars, SEM. Mann-Whitney test, ns: p = 0.0679.

## Movie Legends

**Video S1. Germinal vesicle breakdown triggers blastodisc formation, Yg fusion and Cg outward flow, Related to Figure 1.**

Part I: Time-lapse bright-field (left) and fluorescence (right) movie (42 s interval) of an exemplary Tg(*hsp:clip170-GFP*) oocyte labeling the ooplasm during oocyte maturation. Time: 0 to 280 min after maturation induction. Part II-left: Time-lapse fluorescence movie (157 s interval) of an exemplary Tg(*hsp:clip170-GFP*) oocyte marking the ooplasm (cyan) and exposed to Lysotracker marking yolk granules (magenta) during oocyte maturation. Time: 0 to 270 min. Part II-right: Time-lapse fluorescence movie (133 s interval) of maximum intensity projection of an exemplary *wild type* oocyte exposed to Lysotracker to label yolk granules during oocyte maturation. Time: 0 to 270 min. Part III-left: Time-lapse fluorescence movie (88 s interval) of an exemplary Tg(*hsp:clip170-GFP*) oocyte marking the ooplasm (cyan) and exposed to Lysotracker labeling yolk granules (magenta) during oocyte maturation. Time: 100 to 270 min. Part III-right: Time-lapse fluorescence movie (158 s interval) of an exemplary oocyte injected with Dextran Alexa 647 to label the nucleoplasm of the germinal vesicle during oocyte maturation. Scale bars: 100 μm. Time: 0 to 270 min.

**Video S2. Actin and microtubules are required for ooplasmic reorganization during oocyte maturation, Related to Figure S2.**

Time-lapse fluorescence movie of exemplary Tg(*hsp:clip170-GFP*) oocytes labeling the ooplasm and exposed to DMSO (control, 225 s interval), 100 μM para-Nitroblebbistatin (PBb, 280 s interval), 30 μg/ml Cytochalasin B (Cyto B, 304 s interval), 700 μM Cycloheximide (CHX, 386 s interval, added at 45 minutes after maturation onset) and 200 μM Colchicine (Colchi, 265 s interval) during oocyte maturation. Scale bars: 100 μm. Time: 0 to 270 min after maturation induction.

**Video S3. The actin cytoskeleton rearranges during oocyte maturation, Related to Figure 2.**

Part I: Time-lapse fluorescence movie (182 s interval) of an exemplary Tg(*actb1:Utr-GFP*) oocyte labeling F-actin during oocyte maturation. Time: 0 to 270 min after maturation induction. Part II: Time-lapse fluorescence movie (223 s interval) of an exemplary *wild type* oocyte injected with Dextran Alexa 647 to mark the ooplasm. Time: 0 to 270 min. Part III: Time-lapse fluorescence movie (138 s interval) of an exemplary *wild type* oocyte injected with *CyclinB-GFP* mRNA to visualize CyclinB dynamics. Scale bars: 100 μm. Time: 0 to 270 min.

**Video S4. The bulk actin network reorganization is synchronous with the cell cycle progression, Related to Figure S3.**

Time-lapse fluorescence movie (54 s interval) of an exemplary Tg(*actb1:Utr-mCherry);(Xla.Eef1a1:dclk2a-GFP*) oocyte labeling F-actin (yellow, left) and microtubules (purple, right), respectively, during oocyte maturation. Scale bar: 100 μm. Time: 0 to 270 min after maturation induction.

**Video S5. Microtubule network forms asters upon germinal vesicle breakdown, Related to Figure 3.**

Part I: Time-lapse fluorescence movie (29 s interval) of an exemplary Tg(*Xla.Eef1a1:dclk2a-GFP*) oocyte labeling microtubules during oocyte maturation. Scale bar: 100 μm. Time: 0 to 270 min after maturation induction. Part II: Maximum intensity projection high-resolution time-lapse fluorescence movie (52 s interval) of an exemplary Tg(*Xla.Eef1a1:dclk2a-GFP*) oocyte marking microtubules during oocyte maturation. Scale bar: 50 μm. Time: 0 to 270 min.

**Video S6. Microtubule network transformation is cell cycle-dependent, Related to Figure 3.**

Time-lapse fluorescence movie of exemplary Tg(*Xla.Eef1a1:dclk2a-GFP*) oocytes labeling microtubules and exposed to DMSO (left, 159 s interval, added at 95 min), Cycloheximide (CHX, middle, 204 s interval, added at 85 min) or Dinaciclib (right, 165 s interval, added at 38 min) during oocyte maturation. Scale bars: 100 μm. Time: 0 to 270 min after maturation induction.

**Video S7. Microtubule aster formation relies on dynein activity, Related to Figure 4.**

Time-lapse fluorescence movie of exemplary Tg(*Xla.Eef1a1:dclk2a-GFP*) oocytes labeling microtubules and exposed to DMSO (top-left, 126 s interval), 300 μM Colchicine (Colchi) without DHP (immature oocytes, top-middle, 145 s interval), 75 μM Ciliobrevin D (Cili D, top-right, 225 s interval, undergoing large-scale contraction), 75 μM Cili D (bottom-left, 225 s interval, forming abnormal asters), 12.5 μM Taxol (bottom-middle, 229 s interval) or 50 μM Taxol (bottom-right, 185 s interval) during oocyte maturation (in the presence of DHP). Scale bars: 100 μm. Time: 0 to the end time point of contractions for each case.

**Video S8. Partial microtubule depolymerization drives microtubule aster formation during oocyte maturation, Related to Figure 4.**

Part I: Time-lapse fluorescence movie (275 s interval) of an exemplary Tg(*Xla.Eef1a1:dclk2a-GFP*) oocyte labeling microtubules (left) and injected with Dextran Alexa 647 to mark the nucleoplasm of the germinal vesicle (right) during oocyte maturation. Time: 0 to 270 min after maturation onset. Part II: Time-lapse fluorescence movie of exemplary immature Tg(*Xla.Eef1a1:dclk2a-GFP*) oocytes (in the absence of DHP) labeling microtubules exposed to DMSO (control, left, 161 s interval, added at 72 min) or 300 μM Colchicine (Colchi, right, 144 s interval, added at 67 min). Time: 0 to 456 min after incubation. Scale bars: 100 μm.

**Video S9. Microtubule asters are required for chorion elevation upon activation, Related to Figure 5.**

Part I-left: Time-lapse fluorescence movie (110 s interval) of an exemplary Tg(*actb2:Rab11a-NeonGreen*) oocyte tracing Rab11-positive vesicles during oocyte maturation. Scale bar: 100 μm. Time: 0 to 270 min after maturation induction. Part I-right: Time-lapse fluorescence movie (72 s interval) of an exemplary fully-mature Tg(*actb2:Rab11a-NeonGreen*) oocyte marking Rab11-positive vesicles (green) and exposed to Lysotracker to label yolk granules (magenta) during egg activation. Scale bar: 50 μm. Time: 0 to 144 min after exposure to E3 inducing activation. Part II: Time-lapse fluorescence movie (86 s interval) of an exemplary Tg(*actb2:Rab11a-NeonGreen*) oocyte tracing Rab11-positive vesicles (cyan, left) and injected with *DCLK-mKO2* mRNA to label microtubules (magenta, right) during oocyte maturation. Scale bar: 50 μm. Time: 0 to 240 min after maturation induction.

**Video S10. Part I: microtubule aster translocation to the oocyte cortex relies on dynein activity; Part II: blastodisc formation depends on yolk granule fusion, Related to Figures S4 and S6.**

Part I: Time-lapse fluorescence movie of exemplary Tg(*Xla.Eef1a1:dclk2a-GFP*) oocytes labeling microtubules and exposed to DMSO (control, 257 s interval), or 75 μM Ciliobrevin D (Cili D, 225 s interval, forming abnormal asters) during oocyte maturation. Time: 0 to 270 min after maturation induction. Part II: Time-lapse fluorescence movie of exemplary Tg(*hsp:clip170-GFP*) oocytes marking the ooplasm and exposed to DMSO (control, 225 s interval), or 100 μM Ouabain (213 s interval) during oocyte maturation. Scale bars: 100 μm. Time: 0 to 270 min.

**Video S11. Yolk granule fusion is required for cortical granule translocation towards the oocyte cortex, Related to Figures S5 and 6.**

Time-lapse fluorescence movie of exemplary Tg(*hsp:clip170-GFP*) oocytes labeling the ooplasm (cyan) and exposed to Lysotracker, marking yolk granules (magenta), and DMSO (control), 200 μM Colchicine (Colchi), 100 μM para-Nitroblebbistatin (PBb) or 100 μM Ouabain (30 min interval) during oocyte maturation. Scale bars: 100 μm. Time: 0 to 270 min after maturation induction.

**Video S12. K^+^ concentration increase upon germinal vesicle breakdown triggers yolk granule fusion, Related to Figures 6 and S6.**

Part I: Time-lapse fluorescence movie of exemplary *wild type* zebrafish oocytes exposed to Lysotracker, labeling yolk granules, and DMSO (control, left, 164 s interval) or 100 μM Ouabain (right, 108 s interval) during oocyte maturation. Scale bars: 100 μm. Time: 0 to 270 min after maturation induction. Part II: Time-lapse fluorescence movie of exemplary oocytes injected with Dextran Alexa 647 to mark the ooplasm and K^+^ indicator (K^+^ green) during oocyte maturation. Left: *wild type* oocytes (control, 232 s interval). Right: oocytes exposed to 100 μM Ouabain (172 s interval). Scale bars: 50 μm. Time: 0 to 270 min.

## Materials and Methods

### Animal Husbandry

Fish maintenance was carried out as described in (Westerfield, 2000). 6-24 months old females from WT AB, TL and ABXTL strains as well as Tg(*hsp:Clip170-eGFP), Tg(actb1:Utr-GFP), Tg(Xla.Eef1a1:dclk2a-GFP), Tg(actb1:Utr-mCherry);(Xla.Eef1a1:dclk2a-GFP*) and Tg(*actb2:Rab11a-NeonGreen*) zebrafish lines were used in this study. Fish were bred in the zebrafish facility at ISTA according to local regulations, and all procedures were approved by the Ethic Committee of ISTA regulating animal care and usage.

### Ovarian Follicle Isolation

Methods of ovarian follicle isolation and culture were adapted from (Elkouby and Mullins, 2017; Nair et al., 2013). Female fish were anesthetized in 0.02% Tricaine, and euthanized by decapitation. Ovaries were harvested in culture medium [90% Leibovitz’s L-15 medium with L-glutamine (ThermoFisher), pH 9.0, Penicillin-Streptomycin 50 U/ml, and 0.5% bovine serum albumin (BSA, Sigma-Aldrich)]. Follicles were isolated from ovaries by gentle pipetting with a glass Pasteur pipette and dissection with forceps. Stage-III oocytes were kept in culture medium for up to 12 hours at 25-26 °C.

### Transgenic Lines

For live imaging of F-actin and microtubules, oocytes from Tg(*actb1:Utr-GFP*) (actin)*, Tg(Xla.Eef1a1:dclk2a-GFP*) (microtubules) and Tg(*actb1:Utr-mCherry);(Xla.Eef1a1:dclk2a-GFP*) were used (Behrndt et al., 2012; York et al., 2012). Oocytes from Tg(*hsp:Clip170-eGFP*) were used for labeling of the ooplasm. To visualize subcellular Rab11 localization, the Tg(*actb2:Rab11a-NeonGreen*) zebrafish line ubiquitously expressing NeonGreen-tagged Rab11a was generated using the Tol2/Gateway technology (Kwan et al., 2007; Villefranc et al., 2007).

### mRNA Injections

*PCS2-Rab11-mCherry* (Nowak et al., 2011) and *PCS2-Rab11S25N-mCherry* (see below), *PCS2-DCLK-mKO2* (see below), *PCS2-Utrophin-GFP* (Burkel et al., 2007), and *PCS2-CyclinB1-GFP* (Addgene plasmid #128426) expression constructs were used. mRNAs were synthesized using the SP6 mMessage mMachine Kit (ThermoFisher). Injections into stage-III oocytes were performed as described in (Nair et al., 2013) using glass capillary needles (30-0020, Harvard Apparatus, MA, USA), which were pulled by a needle puller (P-97, Sutter Instrument) and attached to a microinjection system (PV820, World Precision Instruments). The *PCS2-Rab11S25N-mCherry* plasmid was generated using Q5 Site-Directed Mutagenesis Kit (NEB) with *PCS2-Rab11-mCherry* plasmid and Rab11S25N-mCherry-Fwd (TGTGGGGAAGaatAACCTGCTGT) and Rab11S25N-mCherry-Rev (CCAGAGTCTCCAATTAGGACC) primers. The *PCS2-DCLK-mKO2* plasmid was constructed by amplifying the coding sequences of *DCLK1a-202-deltaK* (from *pT2KXIG-Xef1a-DCLK-GFP,* ZFIN ID: ZDB-TGCONSTRCT-090702-3) and *mKO2* (from *mKOkappa-2A-mTurquoise2,* Addgene plasmid # 98837) using gene specific primers with overlapping arms DCLK1a-202_FOR (TGCAGGATCCCATATGGAGGAGCATTTTGACGA), DCLK1a-202_REV (taatcacactCCGATCTGAAATGGAGCTC), mKO2_FOR (TCAGATCGGagtgtgattaaaccagagatgaagatga) and mKO2_REV (TCACTATAGTTCTAGAGGCggaatgagctactgcatcttcta) and then cloned into ClaI-HF (NEB) and XhoI-HF (NEB) digested pCS2 plasmid (Lawson #444) with NEBuilder HiFi DNA Assembly Master Mix (NEB). The assembled plasmids were used for the transformation of chemically competent E. coli DH5α cells (NEB) and verified via Sanger sequencing. 350 pg of *Rab11-mCherry,* 350 pg of *Rab11S25N-mCherry*, 450 pg *DCLK-mKO2,* 200 pg *Utrophin-GFP* and 300 pg *CyclinB1-GFP* mRNA were injected into stage-III oocytes. The injected oocytes were incubated in culture medium (see Sample Preparation for Live Imaging section) for 3-4 h prior to maturation induction. The working concentrations of all mRNA solutions were diluted in 0.2M KCl as described (Nair et al., 2013).

### Sample Preparation for Live Imaging

Oocytes were mounted in 0.7% low-melting-point (LMP) agarose (Invitrogen) inside a glass bottom Petri dish (MatTek) and imaged on an inverted Leica SP5 or SP8 confocal microscope equipped with Leica 20x 0.7 NA, 20x 0.75 NA or 40x 1.1 NA objectives or on a spinning disk setup (Andor Revolution Imaging System; Yokogawa CSU-X1) equipped with a Zeiss 40x 1.2 NA water immersion lens (Behrndt et al., 2012) for high-resolution imaging of microtubule asters. Images were rendered and processed using FIJI or Imaris (Bitplane). The temperature during imaging was kept constant at 26 °C using a stage-heating device (Life Imaging services). For drug treatment experiments, Colchicine (Sigma, 200 and 300 μM), Cycloheximide (Sigma, 700 μM), Cytochalasin B (Sigma, 30 μg/ml), Dinaciclib (Selleckchem, 250 μM), DMSO (Sigma), para-Nitroblebbistatin (Optopharma, 100 μM) and Taxol (12.5, 25 and 50 μM, Sigma) were used. For labeling yolk granules, oocytes were treated with Lysotracker Red (Invitrogen, 1 μM). Oocytes were kept in the medium containing drug (and/or Lysotracker) for 1-1.5 h, and then transferred to drug-/Lysotracker-containing LMP agarose for 15-30 min. Prior to start of imaging 6.5 ml culture medium with 1 μg/ml 17alpha,20beta-Dihydroxy-4-pregnen-3-one (DHP, diluted in 100% Ethanol, Sigma) containing drug/Lysotracker was added on top of the solidified LMP agarose. To label the ooplasm or the germinal vesicle (GV) nucleoplasm, 0.5 nl of 2 mg/ml of 10 KDa Dextran Alexa Fluor 647 (Invitrogen) diluted in 0.2 M KCl was injected into the oocyte center or the vicinity of GV, respectively.

### Chorion Elevation Assay

Stage-III oocytes were injected with 350 pg of *Rab11-mCherry* or *Rab11S25N-mCherry* mRNA, incubated in culture medium for 3 h and then induced to undergo maturation by the addition of the DHP hormone for 4.5 h. In the last 30 min of oocyte maturation, the follicle membrane was removed manually using forceps (Nair et al., 2013). The defolliculated fully mature oocytes were then transferred to a dish containing E3 triggering cortical granule exocytosis and thereby chorion elevation (Kanagaraj et al., 2014). After 30 min in the E3 medium, the activated eggs were imaged with a Stereo-microscope and the images were then used to measure the extent of chorion elevation using Fiji software (Schindelin et al., 2012).

### Phalloidin staining

Stage-III oocytes were harvested from *wild type* female fish and induced to undergo maturation by the addition of DHP-containing to the culture medium. After germinal vesicle breakdown (1-1.5 h after maturation onset) the oocytes were fixed in a glass vial containing 2 % paraformaldehyde at 4 °C overnight. Fixed oocytes were then mounted in 4 % LMP agarose and sectioned using a vibratome (VT1200S, Leica) into 200 μm thick slices. Sections were then washed 3 x for 10 min in PBS with 0.1 % Triton X-100, permeabilized by PBS with 0.5 % Triton X-100 for 1 h at room temperature, and blocked in PBS containing 10 % goat serum, 1% DMSO and 0.1% Triton X-100 (blocking buffer) for 3 h at room temperature. For visualizing the bulk actin gradient, the slices were incubated in Phalloidin 488 (Invitrogen, dissolved 1:200 in blocking buffer) at 4 °C overnight and washed 3 x for 10 min in PBS with 0.1 % Triton X-100. The stained sections were then imaged using an inverted Leica SP5 confocal microscope equipped with a Leica 20x objective.

### Yolk Granules and Cortical Granules Segmentation

To measure average yolk granule (Yg) size over time, the Lysotracker-exposed oocytes were imaged using an inverted Leica SP5 or SP8 confocal microscope equipped with a Leica 20x objective. Maximum intensity projection images of Ygs were used to segment Yg (Lysotracker-signal positive) using Ilastik software (Sommer et al., 2011). The segmented images were then analyzed in Fiji to measure the 2D cross-sectional area of Ygs over time. For cortical granules (Cgs) density at the oocyte surface, surface images (30-50 μm beneath the cortex) of Tg(*hsp:Clip170-eGFP*) or Tg(*actb2:Rab11a-NeonGreen*) oocytes exposed to Lysotracker were acquired using an inverted Leica SP5 or Sp8 confocal microscope equipped with a Leica 20x objective. Cgs were identified as ooplasm-as well as Lysotracker-negative granules and segmented using Ilastik software. The cross-sectional area of the imaging plane was also segmented with Ilastik, and the sum of the Cg area present on the imaging plane normalized to the area of the plane was measured as the Cg density at the oocyte surface over time using a custom-designed MATLAB script. To measure the Cg distribution along the oocyte radius, a similar approach was used on images taken at 50-75 μm beneath the cortex. Using the segmented images, oocyte center of mass and radius were determined and the distance of each Cg to the oocyte center was measured and normalized to the oocyte radius. These results were then plotted as histogram with bin size of 0.05 along the normalized radial axis.

### Blastodisc Clearance Analysis

Tg(*hsp:Clip170-eGFP*) oocytes were imaged during oocyte maturation using an inverted Leica SP5 or SP8 confocal microscope equipped with a Leica 20x objective. Oocytes with clear animal-vegetal (AV) orientation were selected for further analysis. A kymograph was acquired along a 80 pix wide line covering the oocyte AV axis. A line was then fitted to the blastodisc-yolk granules interface in the kymograph over time until the end of oocyte maturation. Blastodisc clearance was measured from the tangent of the slope of the fitted line.

### Bulk Actin Gradient Analysis

The phalloidin-labeled and sectioned stage-IV oocytes, which had just undergone germinal vesicle breakdown, were imaged using an inverted Leica SP5 or Sp8 confocal microscope equipped with a Leica 20x objective. Oocytes with clear AV orientation were selected for further analysis. Maximum intensity projection images were used to segment bulk actin-containing ooplasmic pockets with Ilastik. The bulk actin intensity within these pockets was measured in Fiji and averaged over a 350 μm wide window centered along the oocyte AV axis.

### Brightfield Intensity Measurement

To measure brightfield intensity during oocyte maturation, rectangular shaped ROIs were defined within the oocyte center and the averaged intensities over time obtained using Fiji.

### F-actin Intensity Measurement

To measure nuclear F-actin intensity within the germinal vesicle (GV) shortly before and after GV breakdown in Tg(*actb1:Utr-GFP*) oocytes, rectangular shaped ROIs were defined with the GV, and the averaged intensities over time were obtained using Fiji. To measure F-actin intensity within the blastodisc of Tg(*actb1:Utr-GFP*) during oocyte maturation, a kymograph was acquired along a 50 pix-wide line covering the oocyte AV axis. A line scan was then performed at the blastodisc region to measure the Factin intensity within the blastodisc region.

### CyclinB Intensity Measurement

To measure CyclinB intensity within the blastodisc of *CyclinB1-GFP* mRNA injected oocytes during oocyte maturation, a kymograph was acquired using Fiji along a 30 pix-wide line covering the oocyte circumference. A line scan was then performed at the blastodisc region to measure the CyclinB intensity at the blastodisc region.

### Microtubule Intensity Measurement

To measure microtubule intensity within the blastodisc of Tg(*Xla.Eef1a1:dclk2a-GFP*) oocytes during oocyte maturation a kymograph was acquired using Fiji along a 50 pix-wide line covering the oocyte AV axis. A line scan was then performed at the blastodisc region to measure the microtubule intensity at the blastodisc region.

### Ooplasmic Flow Measurement

To measure the ooplasmic flows, oocytes were injected with Dextran Alexa 647 to label ooplasm and imaged with an inverted Leica SP8 confocal microscope equipped with a Leica 20x objective. A kymograph was acquired using Fiji along a 50 pix-wide line covering the oocyte AV axis. Ooplasmic flow speeds were then measured from the slopes of lines fitted to the ooplasm pockets flowing towards the animal pole on the kymograph. The initial positions of those pockets were normalized to the length of the AV axis and utilized for obtaining the ooplasmic flow profile.

### Microtubule Aster Number

Microtubule asters were followed over time using Imaris 9.1.2 (using spot detection algorithm) and their tracking data was imported to MATLAB R2019b to obtain their number over time.

### Microtubule Cluster Size Analysis

The surface of Tg(*Xla.Eef1a1:dclk2a-GFP*) oocytes were imaged with an inverted Leica SP5 or SP8 confocal microscope equipped with a Leica 20x objective. Maximum intensity projection of these images (along the z slices) were utilized to obtain temporal intensity projections from the start to the end of the contractions using the Temporal-Color Code plugin in Fiji. The temporally-projected images were segmented with the Ilastik software, to identify the microtubule structures undergoing contractions. Cluster sizes were then measured by the application of the Voronoi algorithm plugin in Fiji to the segmented images. Size of the first and second biggest clusters were then determined by custom-designed MATLAB script.

### K^+^ Vesicle Number Analysis

Superficial stacks of oocytes injected with 0.5 nl of 0.7 mg/ml ION Potassium Green-2 TMA+ Salt, K^+^ indicator (abcam) and 2 mg/ml of 10 KDa Dextran Alexa Fluor 647 (to label ooplasm) diluted in nuclease-free water were imaged with an inverted Leica SP8 confocal microscope equipped with a Leica 40x objective. K^+^ vesicles were then segmented using Ilastik software and their number was counted over time with a custom-designed MATLAB script.

### Quantification and Statistical Analysis

Statistical analysis was performed using Prism 8 (GraphPad Software). Data are represented as mean ± SEM and analyzed with the Mann-Whitney test. A P value < 0.05 was considered statistically significant. Sample size and P values are mentioned within the figure legends.

## Acknowledgements

We would like to thank Edouard Hannezo and members of the Heisenberg group for fruitful discussions, and the imaging and optics, electron microscopy and zebrafish facilities at IST Austria for their continuous support.

